# Principles of Dopamine Binding to Carbon Surfaces

**DOI:** 10.1101/2021.08.24.457508

**Authors:** Gaurang Khot, Neil Shirtcliffe, Tansu Celikel

## Abstract

Fast Scan Cyclic Voltammetry (FSCV) combined with carbon electrodes is considered as the gold standard method for real-time detection of oxidizable neurotransmitters. The bioinert nature, rapid electron transfer kinetics and long-term stability make carbon an attractive material for probing brain electrochemistry. Herein, we first demonstrate a rapid fabrication process of carbonized nanopipettes and subsequently perform experimental measurements and theoretical simulations to study mechanisms of dopamine binding on carbonized surfaces. To explain the kinetics of dopamine oxidation on carbonized electrodes we adapted the electron-proton transfer model originally developed by Compton and found that the electron-proton transfer model best explains the experimental observations. We further investigated the electron-proton transfer theory by constructing a Density Function Theory (DFT) for visualization of dopamine binding to graphite-like surfaces consisting of heteroatoms. For graphite surfaces that are capped with hydrogen alone, we found that dopamine is oxidized, whereas, on graphite surfaces doped with heteroatoms such as nitrogen and oxygen, we found deprotonation of dopamine along with oxidation thus validating our experimental and theoretical data. These observations provide mechanistic insights into multistep electron transfer during dopamine oxidation on graphite surfaces.

**Graphical abstract:** **A**: Pictorial view of the experimental setup of carbonized electrodes. The application of waveform causes the oxidation of dopamine. **B**. Background subtracted voltammogram of dopamine, wherein the waveform applied is -0.4V to 1.3V and cycled back at -0.4V at 200 V s^-1^ at 10 Hz. **C**: A hotspot showing the oxidation and reduction of dopamine, wherein two distinct redox spots can be seen. The first redox spot can be seen at 0.0V and the second one at 0.5V. Thus showing a multistep electron transfer for dopamine. **D**: A DFT model for dopamine’s interaction with graphite surfaces doped with nitrogen atoms. Oxidation of oxygen (red) can be seen with loss of protons.

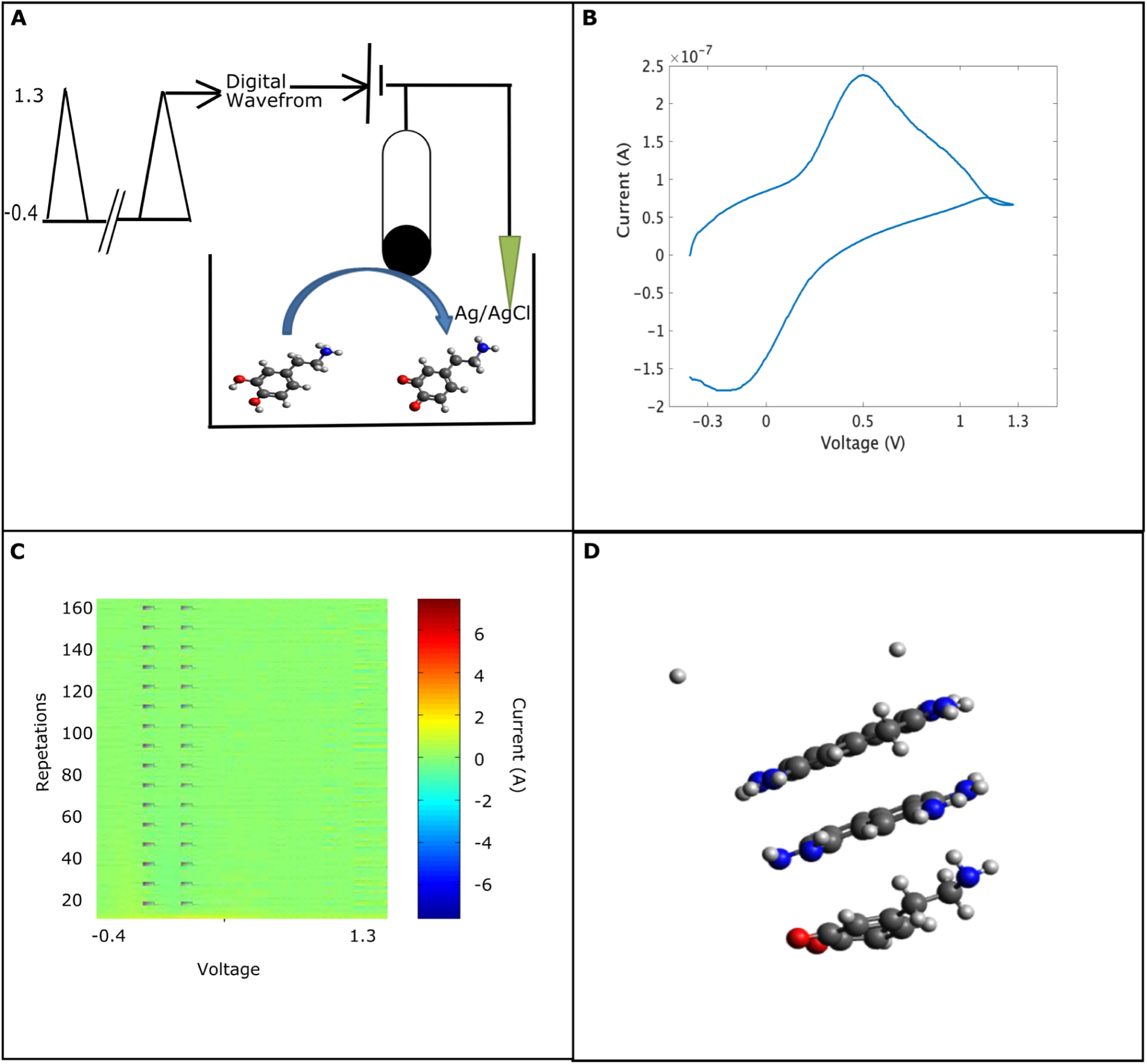

## Introduction

Dopamine is a monoaminergic neurotransmitter that regulates a wide range of neurophysiological functions such as behaviour, attention, cognition (1–4). Low levels of dopamine in the brain are known to cause neurological and psychological disorders like Parkinson’s, Alzheimer’s diseases and schizophrenia (5). Given the widespread roles played by dopamine in neuronal, physiological and cognitive functions, it is important to develop sensors that are able to detect physiological concentrations in the brain.

Methods that are used for *in vivo* detection of dopamine are based on the principles of electrochemistry, spectroscopy or chromatography. Of these methods, electrochemistry has received considerable attention in neuroscience, because of its ability to provide faster sampling time that can match the rapid action potentials thereby allowing to measure the sub-second release of neurotransmitters in the brain (6,7). Another advantage of electrochemistry is its ability to integrate miniature electrodes which can be implanted into tight biological matrices like the brain for understanding neurochemical dynamics (6). These advantages of electrochemical methods have allowed analytical scientists to measure a variety of molecules (neurotransmitters and neuromodulators) in the brain.

Carbon-based allotropes, like graphite, carbon nanotubes, diamonds doped with metals and graphene (8), are commonly used as conductors because of their bio-inertness, stable electrochemical kinetics and low overpotentials compared with metal electrodes (9). Graphite-based materials like carbon fibre electrodes are considered gold standard material for brain electrochemistry because of their biostability and their ability to fabricate microelectrodes that can be implanted in the brain. These microelectrodes have been widely used *in vivo*, *in vitro* and *in silico* for determining neurotransmitters and neuromodulators (10–12).

The sensitivity and rate of the electron transfer process can be improved by means of covalent modifications, which functionalizes the electrode surface (13,14), as well as by the introduction of heteroatoms by performing oxidation of carbon fibre electrodes (15). While considerable progress has been made using carbon fiber electrodes to study chemical neurotransmission, long-term *in vivo* study of neurotransmission is yet to be achieved due to various drawbacks, including 1) the loss of sensitivity occurs which affects the electron transport kinetics of the analyte being measured (16–18). 2) probe degradation, limiting the duration of implants (19,20). Thus there exists a need to develop material that can be used as a long-term implant for dopamine detection.

Pyrolysis of carbon in nitrogen environments offers the possibility of fabricating a graphitic material that can be used as microelectrodes for dopamine detection (21). The advantage of fabrication of electrodes by means of pyrolysis is, it allows rapid fabrication of mechanically strong electrodes whose geometry can be controlled by controlling the pyrolysis process (22–24). Prior research has successfully used pyrolytic electrodes for dopamine detection (21,22,25), however little is understood about the electrochemical kinetics on such a surface. In this report, we expand our understanding of the dopamine oxidation process using Fast scan cyclic voltammetry (FSCV). We fabricate graphite-like carbon microelectrode in a nitrogen-rich environment to study the electrochemical kinetics of dopamine. We hypothesize that introducing heteroatoms on the pyrolytic carbon would promote adsorption, thereby improving the electron and proton transfer process (26,27). To quantify our experimental and theoretical data we construct Density Function Theory (DFT) model (28) to visualize dopamine binding to graphite sheets.

## Materials and Methods

All chemicals were of analytical qualities and were purchased from Sigma Aldrich, unless otherwise stated: Dopamine hydrochloride (DA-HCl), ascorbic acid, 4,-(2-hydroxyethyl-)1-piperazineethanesulfonic acid (HEPES), sodium chloride (NaCl), potassium chloride (KCl), sodium bicarbonate (NaHCO_3_), magnesium chloride (MgCl_2_), monosodium phosphate (NaH_2_PO_4_). Quartz capillaries (o.d. 1 mm, i.d. 0.5mm, length 7.5 cm) were purchased from Sutter Instruments. Deionised (Millipore, 18MΩ) water was used as a solvent.

### 1. Preparation of ultramicroelectrodes

Quartz capillaries (O.D: 1.0mm and I.D: 0.5mm) were pulled using a Sutter puller P-2000 (Sutter instruments). The pulling parameters were: heating temperature 750°C, velocity parameter (apparatus parameter) 50, which gave a tip diameter of 0.8-1.5µm. The tip length was manually reduced to reach a final outer diameter of 25-30µm.

### 2. Fabrication of graphite electrodes via pyrolysis

Pyrolysis was carried out in a counterflow of nitrogen to limit oxidation as described elsewhere (22). Propane was used as the carbon source, the propane tube was connected to the wider end of the capillary, while the immediate vicinity of the narrow end was shielded with nitrogen delivered via a wider quartz tube (Supplementary Figure 1). The heat was delivered to the capillary using a propane-butane torch. The narrow end of the capillary was heated first and the heat was gradually shifted inwards to deposit carbon inside the capillary for electrical connections. Pyrolysis was carried out for 60 seconds. Following the pyrolysis process, electrodes were allowed to cool in nitrogen flow for 2 minutes to limit reaction with air.

### 3. Scanning Electron Microscopy of Electrodes

Scanning electron microscopy was performed using a JEOL 6330 Cryo FESEM. Carbon-filled tips were cut into 2 cm lengths and coated with gold. Imaging was performed at low current to prevent charging.

### 4. Electrochemical setup

The electrochemical setup consisted of a 2 electrode system connected to a patch-clamp amplifier (Intan Technologies, Los Angeles, USA). Briefly, a silver chloride coated silver in KCl solution (3.5M) was used as both reference and counter electrode in a chamber with solvent input and output ports, similar to chambers that are used for electrophysiological recordings in brain slices (29–31). The entire assembly was placed in a Faraday cage to shield it from electrical noise. The experimental setup consisted of a flow injection apparatus where buffer and target analyte were delivered using a peristaltic pump at 32°C. Buffer was delivered at a constant flow rate of 2mL min^-1^ and analyte was introduced in a separate channel at the rate of 1mL min^-1^ for 5 s (total volume ∼100 µL) (16). The electrochemical electrode was positioned near the outlet of the flow of analytes to maximize the response. Electrochemical detection was carried out using a “dopamine waveform” (15). This waveform involved scanning the working electrode from –0.4V to +1.3V and cycling back to -0.4V at a scan rate of 200V s^-1^ repeated at 10Hz, with a step size of 5mV. To obtain a stable background current the electrodes were cycled in a buffer for 30-45 min. at a scan rate of 1000V s^-1^ to precondition them (32). The fast scanning rate resulted in a large capacitive background current that was subtracted digitally using functions written in MATLAB (MathWorks, USA) to extract the true Faradaic current (33). Electrochemical measurement of dopamine was carried in artificial cerebrospinal fluid (aCSF), (31,34,35), consisting of 135mM NaCl, 5.4mM KCl, 5mM Na-HEPES buffer, 1.8mM CaCl_2_ and 1mM of MgCl.

### 5. Construction of Computer Models

To generate input files for ORCA (.inp), Avogadro’s workbench was used wherein models were designed in Avogadro (36) and the source code was implemented in Avogadro (28,37). Geometry optimization was carried out using (!B3LYP-DFT) basis set (28).

#### i. Construction of Graphite Structure Models

Graphite model structures used here were either a stack of 3 layers of carbon atoms or a single carbon monolayer, where the ends were capped with hydrogens or functional groups (**Supplementary information 1-2**). The geometry optimization ensures that a rigid structure is maintained through the simulations thereby preventing the bending of sheets. The number of atoms in the graphite was limited to 36 to facilitate computing efficiency. Graphite interactions occur with one another by means of π-π interactions. The initial placement of molecules resulted in no global minimum and the potential energy was set to 0 (dE=0). To optimize the geometry run, DFT calculations were performed in ORCA, 4.0 (28,37), using an implicit hydration model. In order to reduce noise, the numerical Hessian frequency was used with a cutoff of 50 a.u.. The maximum number of iterations for SO-SCF convergence was set to 124. If the geometry convergence failed within 124 iterations, the structure was discarded and appropriate changes were made for geometry optimization.

#### ii. Construction of Dopamine Structures

Dopamine models were constructed in Avogadro (36) and their geometry optimization was performed in ORCA using DFT model at 310 K using an implicit hydration model (38) (**Supplementary information 3**).

#### iii. Optimization of Transition State for Dopamine and Graphite Surfaces

To simulate the binding of dopamine onto graphite surfaces, a transition search was carried out by connecting dopamine to graphite by means of dummy atoms. The two hydroxyl groups of dopamine were bonded to hydrogen atoms on graphite with no constraints for the bond, or distances (**Supplementary Information 4**). Aqueous medium used as a dielectric constant in our simulations to mimic the interaction of dopamine binding with water. Importantly this only allows for dipole interaction and does not account for hydrogen bonding. After the end of the transition search, bond distance and Gibbs free energy was given by (39,40)

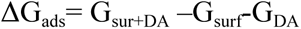

## Results and Discussion

### 1. Fabrication of Carbonization Electrodes

Graphite-like carbonized electrodes were fabricated by pyrolysis of propane at 1bar of atmospheric pressure under a steady stream of nitrogen supplied at 50mL min^-1^ in the opposite direction. Steady heat was delivered to the tip of the carbon nano-pipette by means of a propane-butane torch. The duration of the heating had an effect on carbon deposition on the walls of the quartz capillary. The heat was delivered for time intervals of 20 seconds, 40 seconds and 60 seconds that resulted in decreasing pore opening sizes from ∼25 µm, ∼15 µm and ∼5 µm respectively. The heat was first delivered at the tip of the electrode for a time span of 60 seconds. The delivery of heat was proportional to carbon formation on the edges of the tip. A lower pressure of propane (50-200 millibars) did produce carbonized electrodes, however, the resulting carbon formed weak coupling and disintegrated upon mechanical pressure, e.g. a silver wire was inserted for electrical contact. In order to obtain mechanically more stable carbonized electrodes, we used 1 bar of propane pressure throughout the experiments. To investigate whether the nitrogen flow rate can control the geometry at the tip of the electrode, we manipulated the nitrogen flow rate from 30 mL min^-1^ to 50 mL min^-1^. When a lower flow rate (30 mL min^-1^) was used a cylindrical geometry is obtained at the tip of the electrode (Figure 1A **and** 1B). This resulted in an increase in the surface area and outer diameter of the electrode. Upon increasing the nitrogen flow rate to (50 mL min^-1^) we were able to fabricate electrodes that had circular tips and this was suitable with our electrochemical setup (Figure 1C **and** 1D).

**Figure 1.**
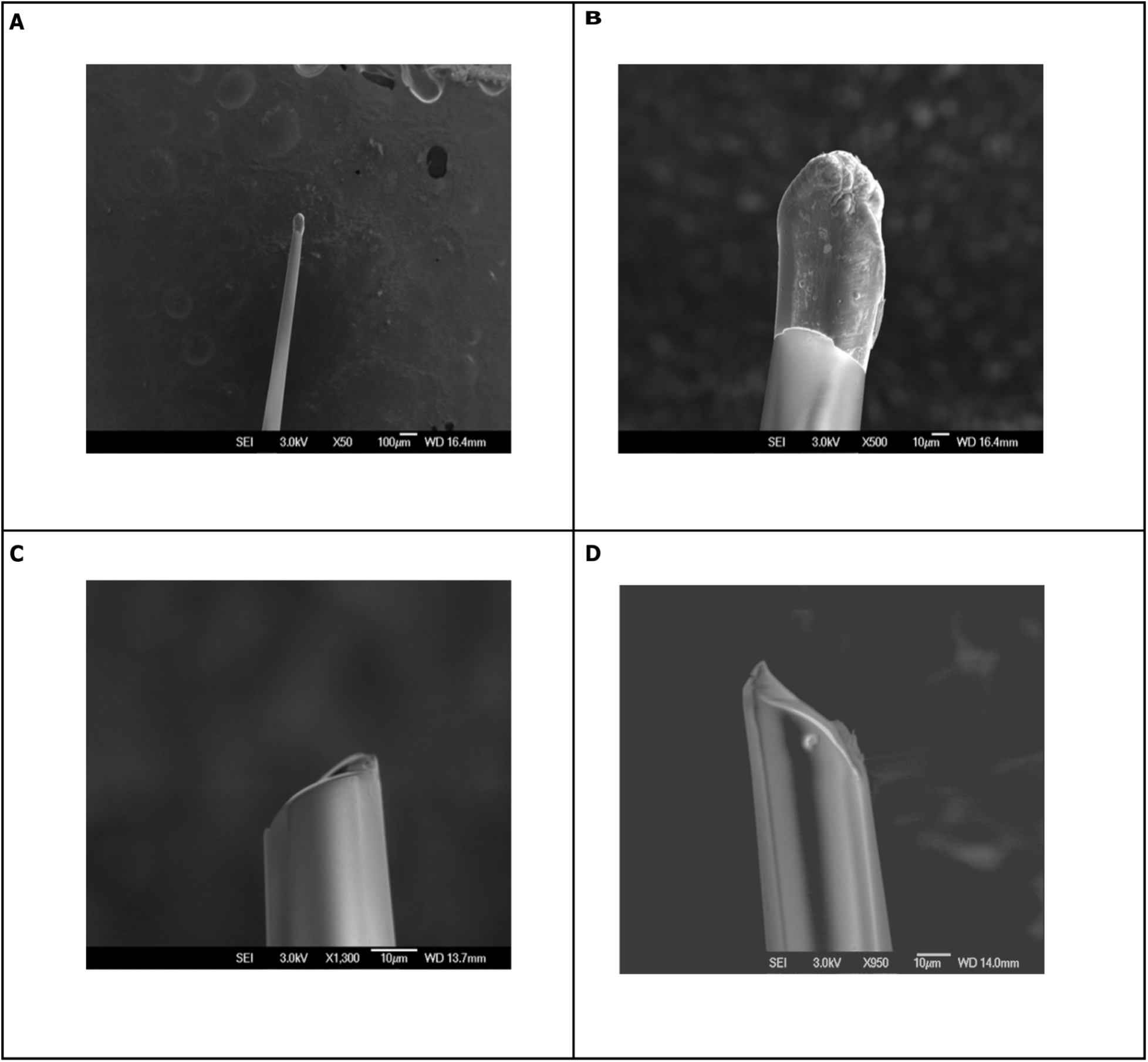
The electron micrograph of carbonized electrodes. **A:** When the flow of nitrogen was kept constant a 30 mL min^-1^ while propane supplied at 1 bar. With a 30-second heating duration a cylindrical electrode that has a radius of 6 μm, a height of 4.5 μm and a surface area of 395.84 μm^2^ was created. Note that because of uneven breakage of the capillary on the formation of the tube the deposition of carbon remains uneven. **B:** Shows the cylindrical electrodes that have a bulb formation at the tip of the electrode. **C:** With the heating duration of 60 seconds and nitrogen flow rate of 50 mL min^-1^ a thin film of carbon is deposited around the walls of the quartz capillaries. This gives a circular electrode having a diameter of 5μm having an area of 19.63μm^2^. **D:** Shows the side view wherein a thin film of carbon can be seen on the walls of capillaries.

### 2. Electrochemical Detection of Dopamine at Pyrolyzed Electrode using FSCV

To study the oxidation of dopamine on carbonized electrodes, 1*μ*M of dopamine was introduced to the flow cell apparatus and a constant flow of buffer was maintained at 2mL min^-1^. Carbonized electrodes having circular geometry were used for understanding dopamine dynamics onto pyrolytic electrodes. To reveal the Faradaic signature of dopamine, background subtraction was performed.

Dopamine shows a broad oxidation and reduction peak at 0.51V and -0.10V respectively (Figure 2A) when the voltage slope was constant at 200V s^-1^ and stimulus was repeated at 10Hz. To understand the transport kinetics of dopamine from bulk to the surface of the electrode, the scan rate was varied and the peak current was plotted against scan rate and the square root of scan rate. As scan rate was proportional to peak current (Figure 2C) adsorption was the mode of transport for dopamine from the bulk solution to the surface of the electrode (41). The electrodes had a sensitivity 0.35 (±0.7) µA *µ*M^-1^ to 1*µ*M of dopamine. The square root of the scan rate was proportional to peak current (Figure 2D). This suggests that dopamine is predominantly adsorbed on the surface of the electrode and once all the adsorption sites are occupied, dopamine from bulk solution diffuses on the surface of the electrode. The plot of the log of peak current against the square root of scan rate shows (Figure 2E) that as the scan rate is increased the electrochemical kinetics improve and dopamine oxidation is quasi-reversible.

**Figure 2.**
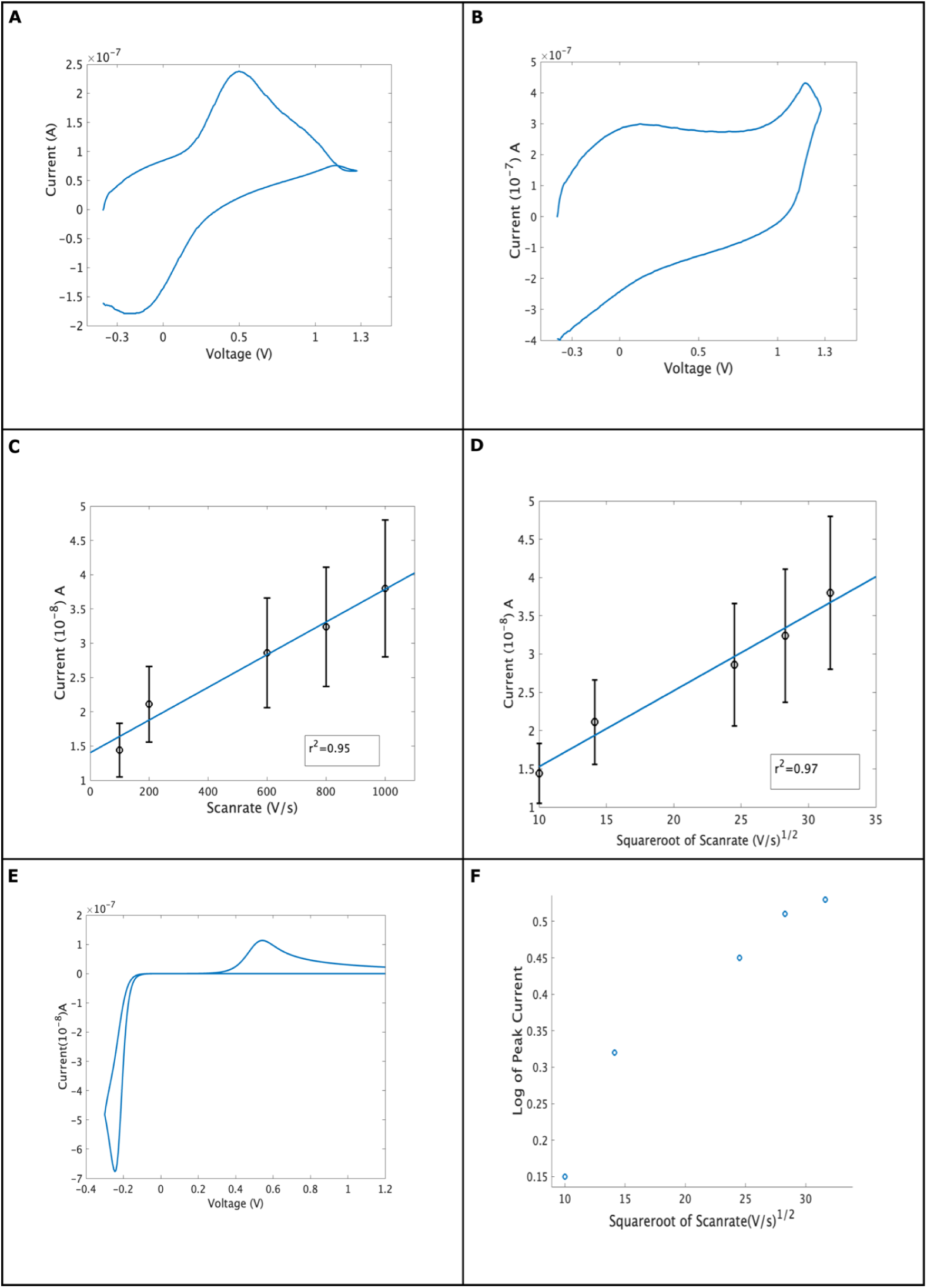
The response of electrode to 1 *µ*M dopamine, wherein the electrode was scanned from -0.4V to 1.3V and cycled back to -0.4V at 200 V s^-1^ at 10 Hz. **A:** The background-subtracted voltammogram for 1 *µ*M of dopamine, on pyrolyzed carbon. Dopamine is oxidized at 0.51V(±0.12V) and reduced at -0.11V(±0.10V). **B:** The resultant background current. **C:** The oxidation peak current increases with scan rate. **D:** Square root of scan rate against peak current. **E:** Simulated peak for dopamine, where dopamine becomes oxidized at 0.57V and reduced at -0.2V. The parameters were, diffusion coefficient of dopamine, 7.6 × 10^-6^ cm^2^ s^-1^, area of the electrode is 19.63μm^2^,(±2.4μm^2^) rate of electron transfer, 3×10^-4^ cm^2^ s^-1^ (26,27). The difference in the magnitude of the experimental and simulated current is because of the roughness in the surface area of the experimental electrode. **F)** the square root of scan rate against peak current.

To understand whether our pyrolytic electrode offers a reduction in potential, we simulated the Randles-Sevick equation using the standard parameters (16). Our simulation result shows that dopamine is oxidized at 0.6V and reduced at -0.1V. A likely reason for the difference in potential from theory to experimental can be attributed to the faster electrochemical kinetics offered by pyrolytic electrodes.

Pyrolytic electrodes are known to reduce the anodic potential for dopamine. It was previously shown that dopamine oxidizes 0.45V and reduction at -0.2V (25). Even though carbon fiber electrodes reported oxidation of dopamine at 0.6V and reduction at -0.4V using a waveform of scan rate of 400V s^-1^ repeated at 10Hz (15). A likely reason for this reduction in the anodic potential is the circular geometry of carbon nanofiber improving the electrochemical kinetics by accelerating the electron transfer process (25). To understand if the difference in oxidation voltages of dopamine on carbon fiber electrodes as compared to ours is purely on scan speed, we increased the scan speed to 600V s^-1^ at 10 Hz (Supplementary Figure 2). Dopamine oxidation is seen at 0.55V and reduction is seen at -0.2V, thus suggesting that surface chemistry produced during pyrolysis consists of heteroatoms that tend to alter the electron transfer kinetics.

In our pyrolytic electrode, the likely composition of heteroatoms is oxygen, nitrogen and other reactive species that are formed by partial pyrolysis of propane under a nitrogen environment. In our experiments, the oxidation of dopamine is seen at 0.51V (Figure 2A) while the simulated cyclic voltammogram using 2 electron transfer process (Figure 2F) shows oxidation at 0.6V. A likely reason for such a reaction to occur on our surface is because nitrogen atoms that are present on the surface of the carbon can act as a base (NH_2_), thereby they can undergo protonation (NH_3_^+^) and deprotonation. This would explain why nitrogen-containing surfaces both increase the current and decrease the overpotential of dopamine oxidation.

### 3. Understanding Dopamine Electrode Dynamics

To understand the kinetics of dopamine on pyrolyzed electrodes, we studied the voltage differences of the oxidation and reduction peaks (Figure 3A). As the scan rate was increased the peak current increased with a shift in oxidation voltage to anodic potential (Figure 2C). The difference between oxidation and reduction voltage for dopamine was calculated and our results show that the difference is greater than 57 mV. This suggests that dopamine oxidation on pyrolytic electrodes is quasi-reversible (Figure 3A). The quasi-reversible kinetics of dopamine must consist of another mechanism apart from electron transfer. Recently it was proposed that the oxidation process of dopamine consists of a multistep process involving proton and electron transfer (26,27). The oxidation scheme for dopamine consists of, dopamine oxidizing from solution, having a rate constant K_1_ serving as a rate-limiting step towards adsorption of dopamine on the surface of the electrode (6,26,27,42,43). Dopamine on the electrode surface undergoes oxidation of 2e^-^ to dopamine-o-quinone on the surface of the electrode, which is brought into the bulk solution during desorption by K_2_ to give dopamine-o-quinone (41,42,44).

**Figure 3.**
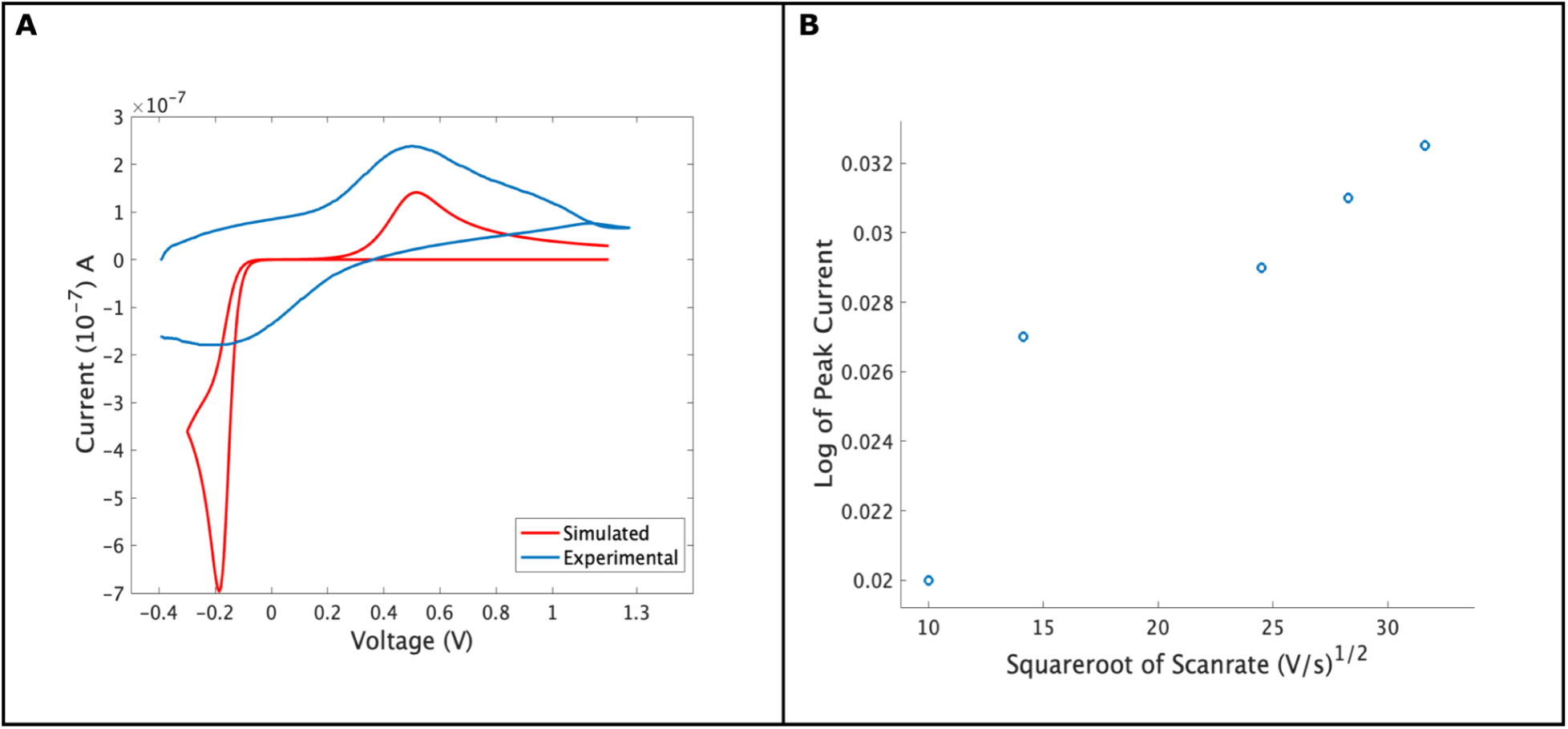
1 *µ*M dopamine’s response to dopamine against different scan speeds. **A:** The difference between oxidation and reduction peaks of dopamine across the square root of scan rate. **B:** Experimental data is matched with simulated data by using electron-proton model (26,27). The parameters for simulations were as follows. The diffusion constant D for dopamine was taken as 7.6 × 10^-6^ cm^2^ s^-1^, the area of the electrode was 0.0001 cm, α=1. The number of electrons transferred was 2, the rate of electron transfer (k) was 0.000003cm/s, the scan speed was 200 V s^-1^ and the temperature was 310 K. It must be noted that the geometric area of the cylindrical electrode was 19.27 μm^2^.

**Figure 4.**
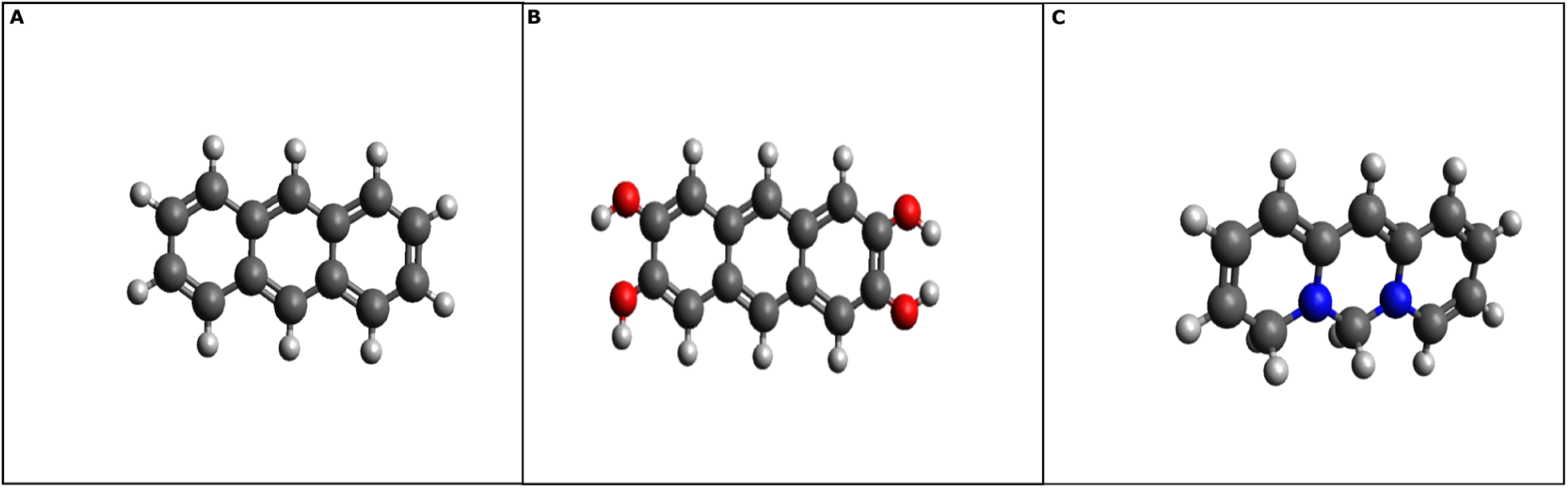
**A:** A model graphite surface. **B:** Carbon surface doped with nitrogen atoms. It should be noted that nitrogen here has a partial positive charge and the structure is for simulation only. **C:** terminal hydroxyl groups.

While recent evidence suggests that a multi-step process occurs wherein the first electron transfer process is the rate-limiting step followed by proton transfers that are rate-determining steps. The second electron transfer occurs to complete the oxidation mechanism, wherein dopamine is oxidized from dopamine to dopamine-o-quinone structure (27). Such a mechanism must be able to accelerate the electron transfer process thus allowing experimental data to match with theory. We initially conducted a simulation of the Butler-Volmer equation using the two-electron transfer kinetics data (Figure 2F) wherein the rate of electron transfer process was set at 3X10^-4^ cm^2^ s^-1^ (26) and found that dopamine oxidation is seen at 0.6V and reduction is seen at -0.1V. Upon adapting the reversibility parameters from the electron-proton model to 10^-4^ m s^-1^ and setting the rate of electron transfer to 3X10^-6^ cm^2^ s^-1^ (16) we see that experimental data match with theory (Figure 3B) (26,27). A considerable difference exists between the experimental and simulation part, as the model does not account for the surface roughness.

FSCV protocols allow superior temporal resolution and a faster sampling rate (6). The faster scan speeds employed to detect dopamine at a subsecond time frame often limit the chemical reactions occurring at the surface of the electrode (6). In our experiments adsorption was the predominant mode of transport for dopamine from bulk to the surface of the electrode (Figure 2C). The adsorption is transient however it is limited, as there are fewer sites for dopamine to adsorb on the surface of the electrode. Once these sites are occupied then the diffusion process dominates the reaction (Figure 2D) wherein time constants are typically in the order of 10^-9^ seconds. Given the high time rate of the diffusion process and the faster rate of electron transfer, it is likely that proton transfer might be hindered. This suggests that graphite-like surfaces must have a higher affinity for electron transport and possibly hinder proton transport.

### 4. Computational Modeling of Dopamine Binding to Graphite Surfaces

To investigate the binding of dopamine onto graphite surfaces, we constructed models of graphite and dopamine separately and optimized their geometry.

#### I. The Model of Graphite Surface

Graphite surfaces consist of carbon atoms that are arranged in sp^2^ hybridization in the form of AB stacking (45,46). The sp^2^ hybridization allows stacking of multiple sheets of carbon atoms via pi-pi bonds and weak Van-der Waals forces (46). In our experiments we used a stack consisting of 3 layers of graphite or anthracene molecules and heteroatoms were added onto edges of the simulated graphite (**Supplementary Information 1, 2**). As we could not obtain the values at conduction and valence band (40eV) (46), we did not consider further thermochemistry run on our model graphite. We optimized the geometry for the dopants and if geometry optimization was not successful hydrogen atoms were placed at the terminal ends to satisfy the valency.

#### II. Dopamine Binding on Carbon Surface

Once we optimized the graphite model structures, we then investigated the effect of dopamine binding onto carbon. At neutral pH, dopamine has an energy level of 4.01eV and an electric dipole moment of 2.94eV. As the pH approaches 3, the energy level jumps to 4.79eV while at basic pH of >13, the energy level is steady at 4.4eV. To understand the electrochemical interactions of graphite occurring with dopamine we connected the two molecules by means of dummy atoms in an aqueous solvent medium (**Supplementary Information 3**). As expected, dopamine lost terminal hydrogen at the start of the simulations and oxygen formed the double bonds (Figure 5A). However, this was seen only for surfaces that had oxygen and nitrogen groups. Graphite surfaces that had only hydrogen atoms tend to form bonds with dopamine, thus showing that hydrogenated surfaces might undergo biofouling by adsorption of dopamine (Figure 5B). Surfaces containing nitrogen, either terminal or doped, rather showed repulsion between dopamine and the surface, thus suggesting a complex electron transfer process (Figure 5C). The bond distance for the surface with oxygen or hydrogen was 2.84 Ångstroms (Å) and 0.8 Å respectively, while it was greater, i.e. 4.84 Å, for nitrogen-containing surfaces. The electrochemical study has shown that different surface chemistry dictates surface adsorption, sensitivity and electrochemical transfer kinetics for dopamine (13). We hypothesize that nitrogen-doped surfaces carry partial charge on the end functional groups and thus they can act as proton receivers while inhibiting binding to the catechol group (Figure 5C).

**Figure 5.**
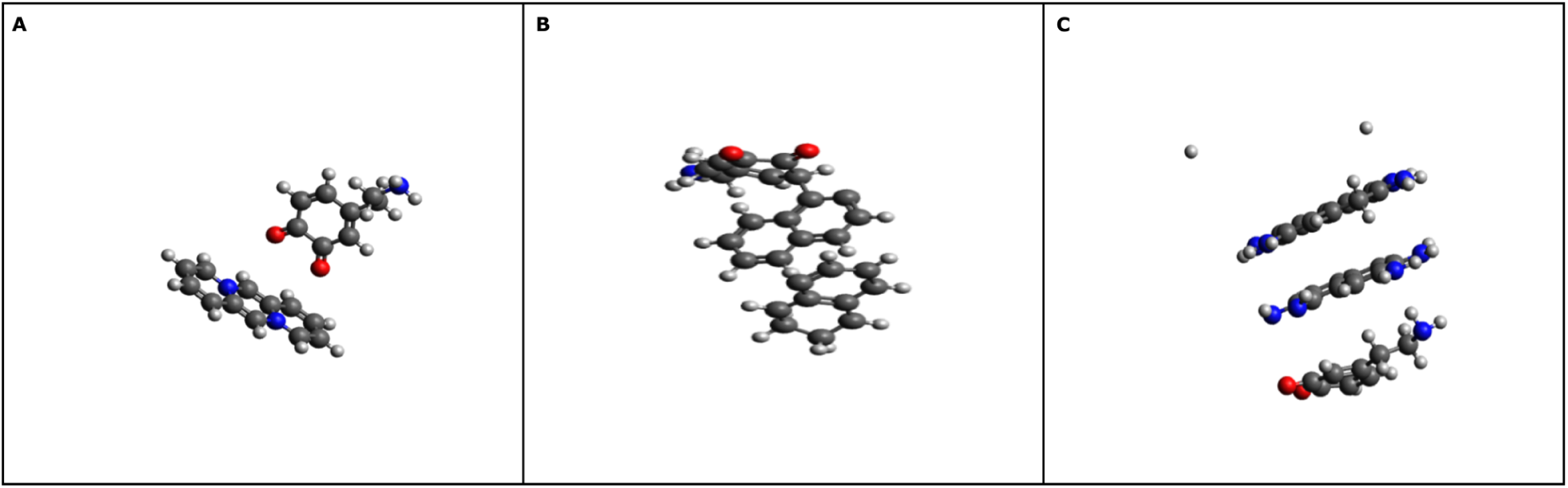
The interaction of dopamine with carbon surfaces on different surface chemistry **A:** The oxidation of dopamine onto nitrogen-doped carbonized surfaces. It can be seen that dopamine oxidizes rapidly when nitrogen atoms are present on the graphite surface at the basal plane. Similar results were obtained for the surface with oxygen functional groups (data not shown). **B:** The binding of dopamine to carbon surfaces. The end group of dopamine is oxidized. **C:** The transfer of protons on carbon surfaces, containing amine end groups. Deprotonation occurs on dopamine structure, suggesting that oxygen groups on dopamine can interact with nitrogen bases on the graphite surface to transfer protons to and from it during redox reactions. The interaction of dopamine with the carbon surface would be expected to occur via pi-pi stacking and by hydrogen bonds.

Our studies make use of an aqueous medium where ionic interactions occur between molecules. When we performed our simulation in a vacuum we found bonding between the lone pairs on the oxygens of dopamine and terminal hydrogens on the edge of the model graphite. Our results were identical with previous work performed that showed the binding of dopamine onto oxygen motif surfaces (39,40). Our goal was to mimic the electrochemical setup and understand whether heteroatoms placed on graphite surfaces behave differently than mono atoms. The adsorption energy for dopamine is on a surface containing terminal nitrogen groups (−6 Kcal), followed by oxygen (−8 Kcal) and then hydrogen (−2.8kcal). Thus showing that dopamine has higher affinity for surfaces with oxygen groups, consistent with electrochemical observations (13–15,40). Limitations of our DFT model to visualize the binding of dopamine to graphite were: 1) Our modelling was limited to a few numbers of atoms for graphite atoms. This resulted in an unrealistic bandgap, thus allowing dopamine to bind readily to graphite atoms. Our binding of graphite to dopamine did not encompass specific geometry or crystalline structure, 2) we made several attempts to define geometry but our convergence failed and thus we opted to carry our simulations in free space with hydration energy. While our current studies have shown the first principle study for dopamine binding to mono-layer of graphite, our future modelling work will take into account the stack of graphite binding to dopamine.

To understand the response of pyrolytic electrodes for detection of dopamine and serotonin, simultaneously, we exposed 1*µ*M of dopamine and serotonin simultaneously(Figure 6A). We find that a single oxidation peak at 0.6V and a reduction peak at 0.0V. To investigate the occurrence of this peak, we exposed the electrode to 1*µ*M of serotonin (Figure 6B). Serotonin is oxidized at 0.8V and reduced at -0.1V. Thus the result shows that pyrolytic electrodes are unable to distinguish the multiplex detection of dopamine and serotonin, because multimodal detection requires surface to alter the potentials for multiple analytes and possibly act as local storage of protons, which is not possible in the case of pyrolytic electrodes.

**Figure 6.**
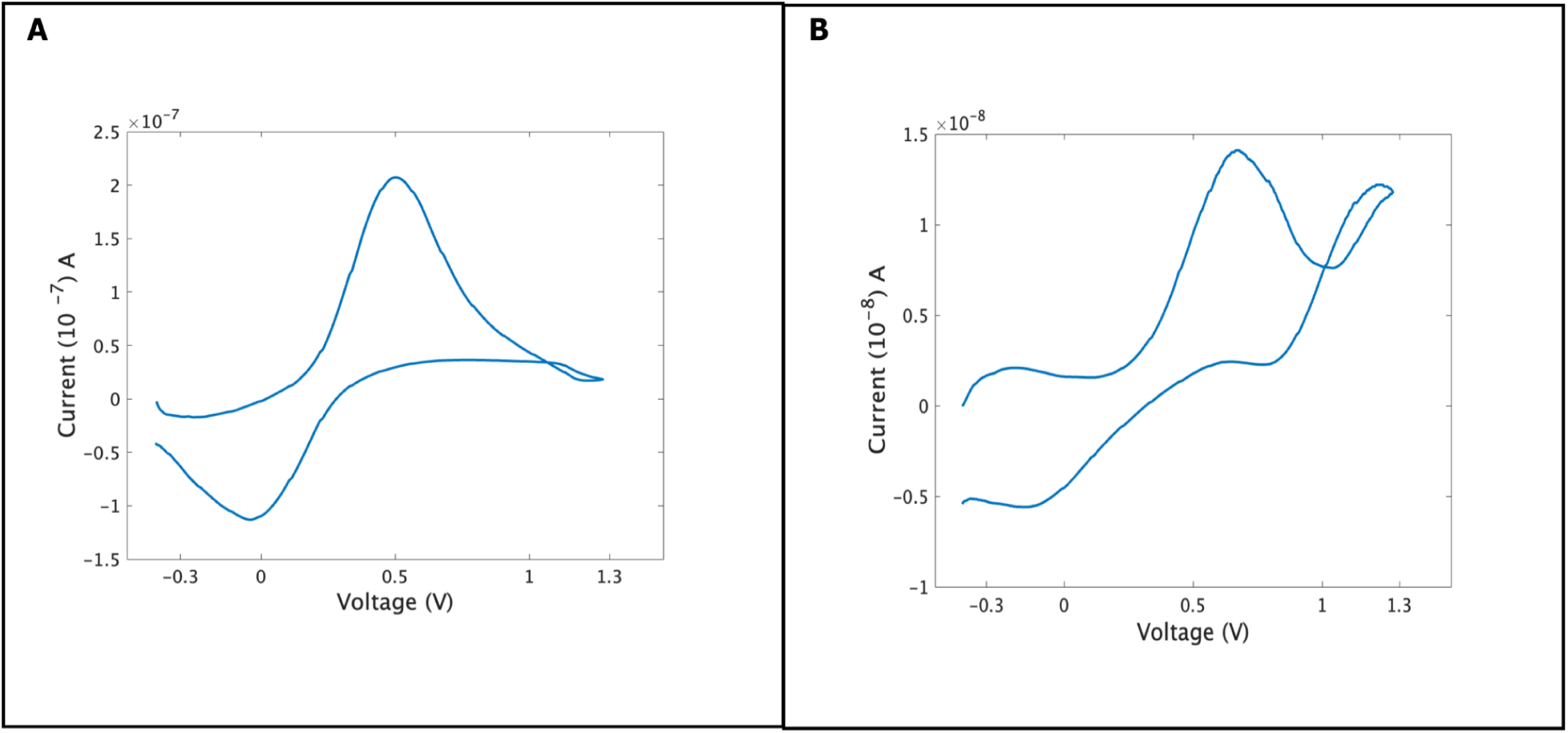
Shows the response of pyrolytic electrodes to dopamine and serotonin. **A:** 1*µ*M of dopamine and serotonin are exposed to pyrolytic electrodes. A single oxidation peak is seen at 0.6V and reduction peak is seen at 0.0V. **B:** 1*µ*M of serotonin is exposed to pyrolytic electrodes. An oxidation peak is seen at -0.2V and a second oxidation peak is seen at 0.8V. A reduction peak can be seen at 1.1V, which is hypothesized by indol-peroxide complex. A second reduction peak is seen at -0.1V which is associated with reduction of serotonin.

Further experimental work would consist of quantification of dopamine and serotonin onto carbonized electrodes for *in vivo* detection and improving the parameters of our DFT models to enhance our understanding of dopamine -graphite interactions.

## Conclusions

In this present report, we fabricated graphite surfaces that are functionalized with nitrogen atoms. The electrochemical reaction kinetics of dopamine were studied on the electrode using FSCV. We found that the electron-proton transfer mechanism explains the quasi-reversible nature of dopamine. To understand the dynamics of dopamine behavior on carbon surfaces, we created a model that explains the molecular kinetics of dopamine onto carbon surfaces, thus confirming our approach on using the electron-proton approach to explain the behavior of dopamine onto carbon surfaces.

## Acknowledgements

This work was supported in part by European Regional Development Fund (MIND, nr. 122035). Author declares no competing interests.

## Supplementary Information

**Supplemental Figure 1.**
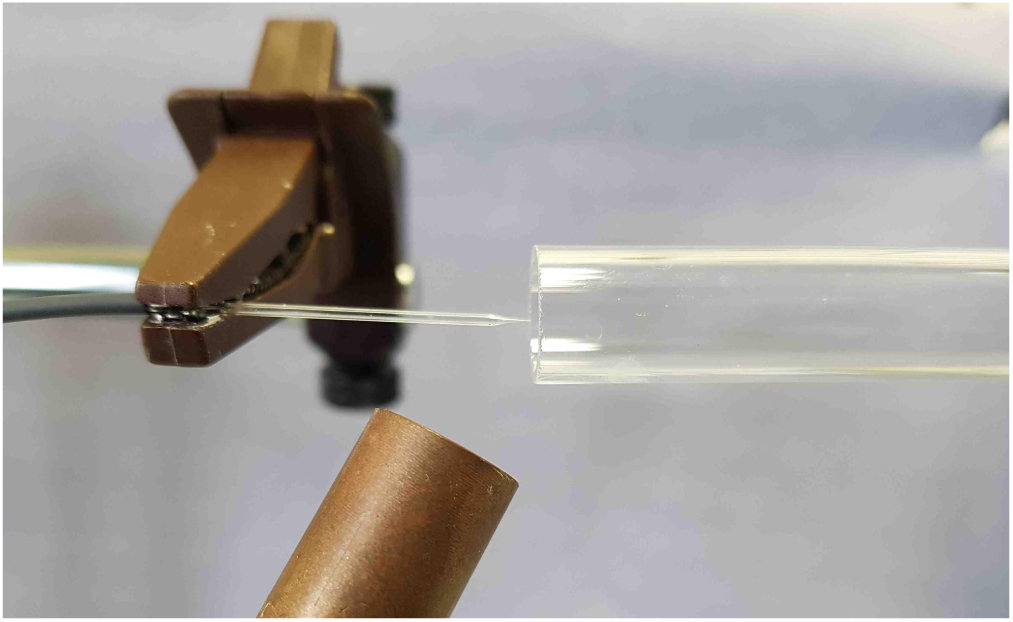
Pyrolysis setup for carbon deposition. The quartz capillary is connected to a low-pressure propane/butane supply at the wider end and a nitrogen shielding gas flow provided via a larger quartz tube (10 mm i.d.). Heat is delivered via propane-butane torch (tip visible at the bottom of this image). Heat is delivered to the narrow end of the capillary and the process of carbonization occurs at the tip.

**Supplemental Figure 2.**
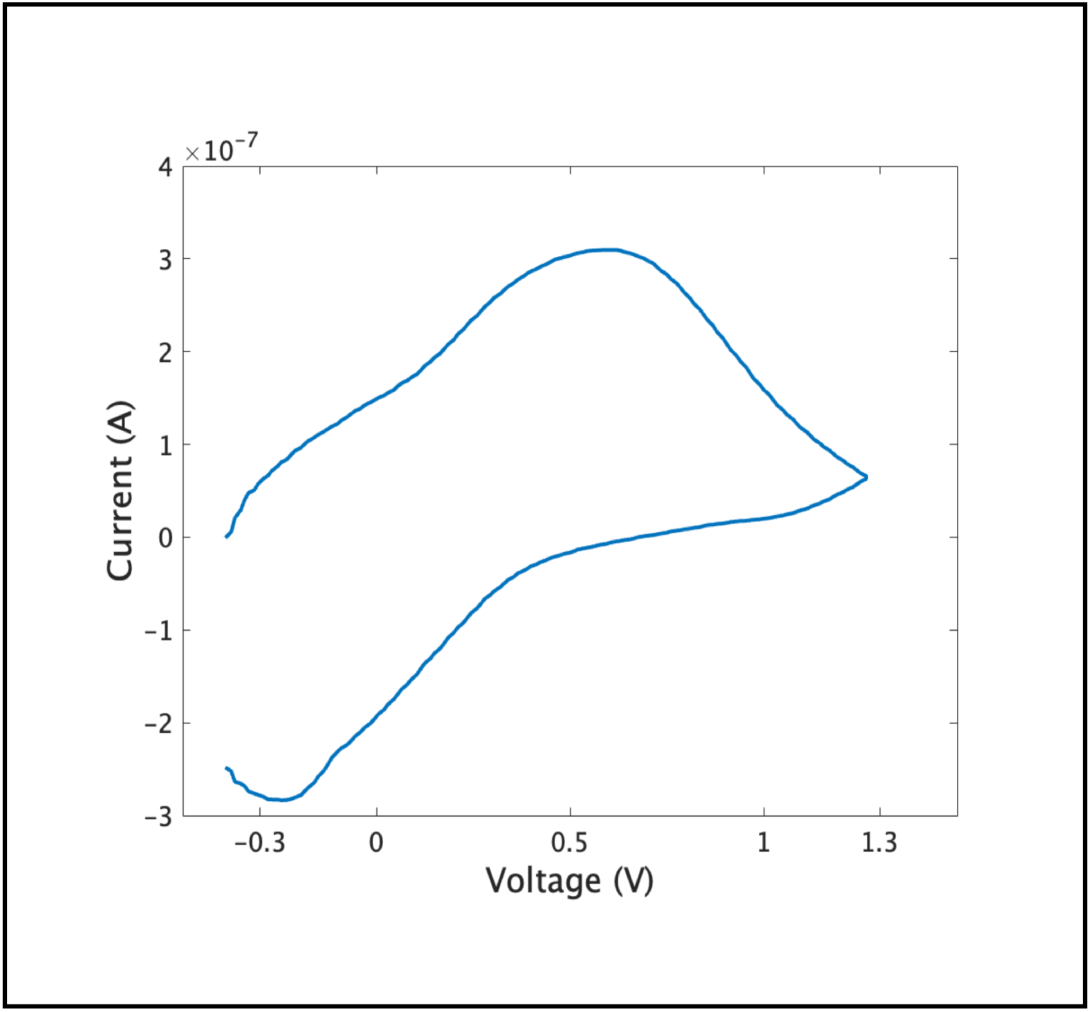
Oxidation of dopamine using the T600 waveform. The waveform parameters were scan speed 600 V s^-1^ from -0.4V to 1.3V and cycled back at -0.4V at 10 Hz. 1*µ*M Dopamine is oxidized at 0.57 V and reduced at -0.23V.

### Supplemental information 1: Molecular input for graphite

|  |  |  |  |
| --- | --- | --- | --- |
| C | 0.00000 | 0.00000 | 0.00000 |
| C | 0.00000 | 0.00000 | 6.79000 |
| C | -1.23500 | 2.13908 | 0.00000 |
| C | -1.23500 | 2.13908 | 6.79000 |
| C | 2.47000 | 0.00000 | 0.00000 |
| C | 2.47000 | 0.00000 | 6.79000 |
| C | 1.23500 | 2.13908 | 0.00000 |
| C | 1.23500 | 2.13908 | 6.79000 |
| C | 0.00000 | 0.00000 | 3.39500 |
| C | -1.23500 | 2.13908 | 3.39500 |
| C | 2.47000 | 0.00000 | 3.39500 |
| C | 1.23500 | 2.13908 | 3.39500 |
| C | 0.00000 | 1.42606 | 0.03395 |
| C | 1.23500 | 0.71303 | 3.42895 |
| C | -1.23500 | -0.71303 | 0.03395 |
| C | 1.23500 | -0.71303 | 0.03395 |
| C | -1.23500 | -0.71303 | 6.82395 |
| C | 1.23500 | -0.71303 | 6.82395 |
| C | 0.00000 | 1.42606 | 6.82395 |
| C | -2.47000 | 1.42606 | 0.03395 |
| C | -1.23500 | 3.56514 | 0.03395 |
| C | -2.47000 | 1.42606 | 6.82395 |
| C | -1.23500 | 3.56514 | 6.82395 |
| C | 3.70500 | -0.71303 | 0.03395 |
| C | 2.47000 | 1.42606 | 0.03395 |
| C | 3.70500 | -0.71303 | 6.82395 |
| C | 2.47000 | 1.42606 | 6.82395 |
| C | 1.23500 | 3.56514 | 0.03395 |
| C | 1.23500 | 3.56514 | 6.82395 |
| C | 0.00000 | -1.42606 | 3.42895 |
| C | -1.23500 | 0.71303 | 3.42895 |
| C | -2.47000 | 2.85211 | 3.42895 |
| C | 0.00000 | 2.85211 | 3.42895 |
| C | 2.47000 | -1.42606 | 3.42895 |
| C | 3.70500 | 0.71303 | 3.42895 |
| C | 2.47000 | 2.85211 | 3.42895 |
| H | -1.23500 | 2.13909 | -1.07000 |
| H | -1.23500 | 2.13909 | 5.72000 |
| H | 2.47000 | -0.00001 | 2.32500 |
| H | -2.04862 | -0.01889 | 0.00090 |
| H | -1.29138 | -1.28597 | 0.93587 |
| H | -1.29138 | -1.36907 | -0.80945 |
| H | 1.23500 | -1.74386 | 0.08303 |
| H | -1.04067 | -1.76471 | 6.79090 |
| H | -1.83134 | -0.43383 | 5.98055 |
| H | -1.75937 | -0.47538 | 7.72587 |
| H | 1.23500 | -1.74386 | 6.87303 |
| H | -2.75800 | 1.25977 | 1.05096 |
| H | -2.35237 | 0.48517 | -0.46183 |
| H | -3.22602 | 1.99837 | -0.46182 |
| H | -1.23500 | 3.89769 | 1.05096 |
| H | -2.10865 | 3.93371 | -0.46182 |
| H | -0.36135 | 3.93371 | -0.46182 |
| H | -2.75800 | 1.25977 | 7.84096 |
| H | -2.35237 | 0.48517 | 6.32817 |
| H | -3.22602 | 1.99837 | 6.32818 |
| H | -1.23500 | 3.89769 | 7.84096 |
| H | -2.10865 | 3.93371 | 6.32818 |
| H | -0.36135 | 3.93371 | 6.32818 |
| H | 4.51862 | -0.01889 | 0.00090 |
| H | 3.76138 | -1.36907 | -0.80945 |
| H | 3.76138 | -1.28597 | 0.93587 |
| H | 3.36273 | 1.94147 | 0.08303 |
| H | 4.51862 | -0.01889 | 6.79090 |
| H | 3.76138 | -1.36907 | 5.98055 |
| H | 3.76138 | -1.28597 | 7.72587 |
| H | 3.36273 | 1.94147 | 6.87303 |
| H | 2.24295 | 3.92269 | 0.00090 |
| H | 0.76701 | 3.90043 | 0.93587 |
| H | 0.69504 | 3.94199 | -0.80945 |
| H | 2.24295 | 3.92269 | 6.79090 |
| H | 0.76701 | 3.90043 | 7.72587 |
| H | 0.69504 | 3.94199 | 5.98055 |
| H | 1.00795 | -1.78361 | 3.39590 |
| H | -0.53996 | -1.80290 | 2.58555 |
| H | -0.46799 | -1.76135 | 4.33087 |
| H | -2.12773 | 0.19761 | 3.47803 |
| H | -3.28362 | 2.15797 | 3.39590 |
| H | -2.52638 | 3.50815 | 2.58555 |
| H | -2.52638 | 3.42505 | 4.33087 |
| H | 0.00000 | 3.88294 | 3.47803 |
| H | 2.47000 | -1.75861 | 4.44596 |
| H | 3.34365 | -1.79463 | 2.93318 |
| H | 1.59635 | -1.79463 | 2.93318 |
| H | 3.99300 | 0.87931 | 4.44596 |
| H | 3.58737 | 1.65392 | 2.93317 |
| H | 4.46102 | 0.14071 | 2.93318 |
| H | 3.28362 | 2.15797 | 3.39590 |
| H | 2.52638 | 3.42505 | 4.33087 |
| H | 2.52638 | 3.50815 | 2.58555 |

\*

### Supplemental Information 2: Molecular input for N-doped graphite

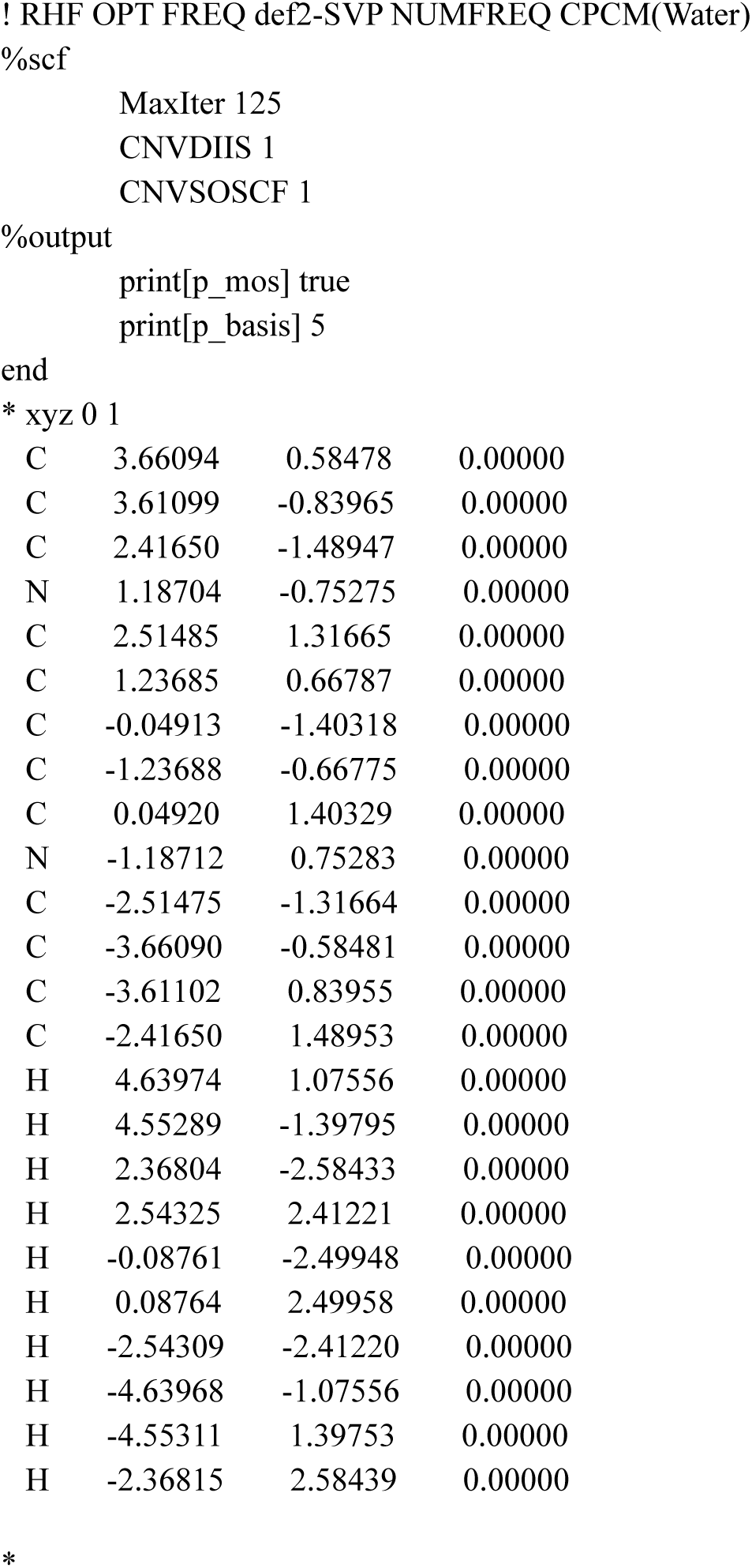

### Supplemental Information 3: Molecular model for oxygen-doped graphite

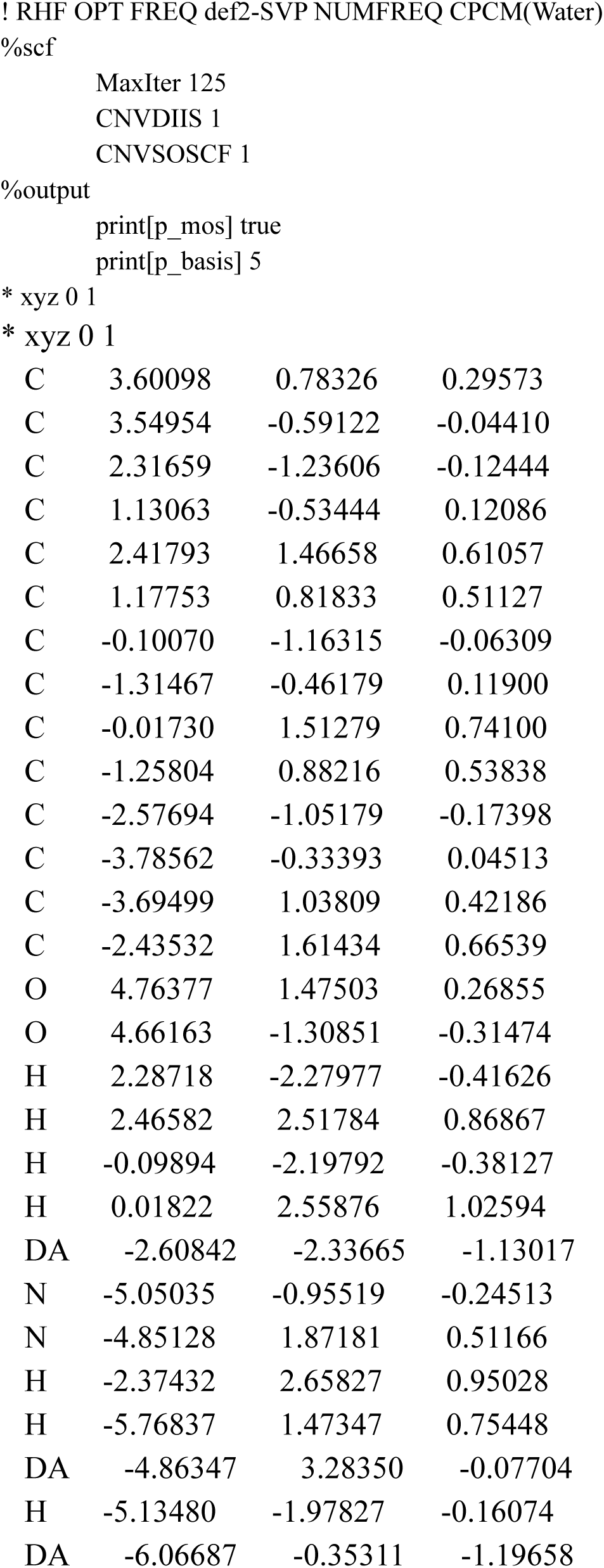
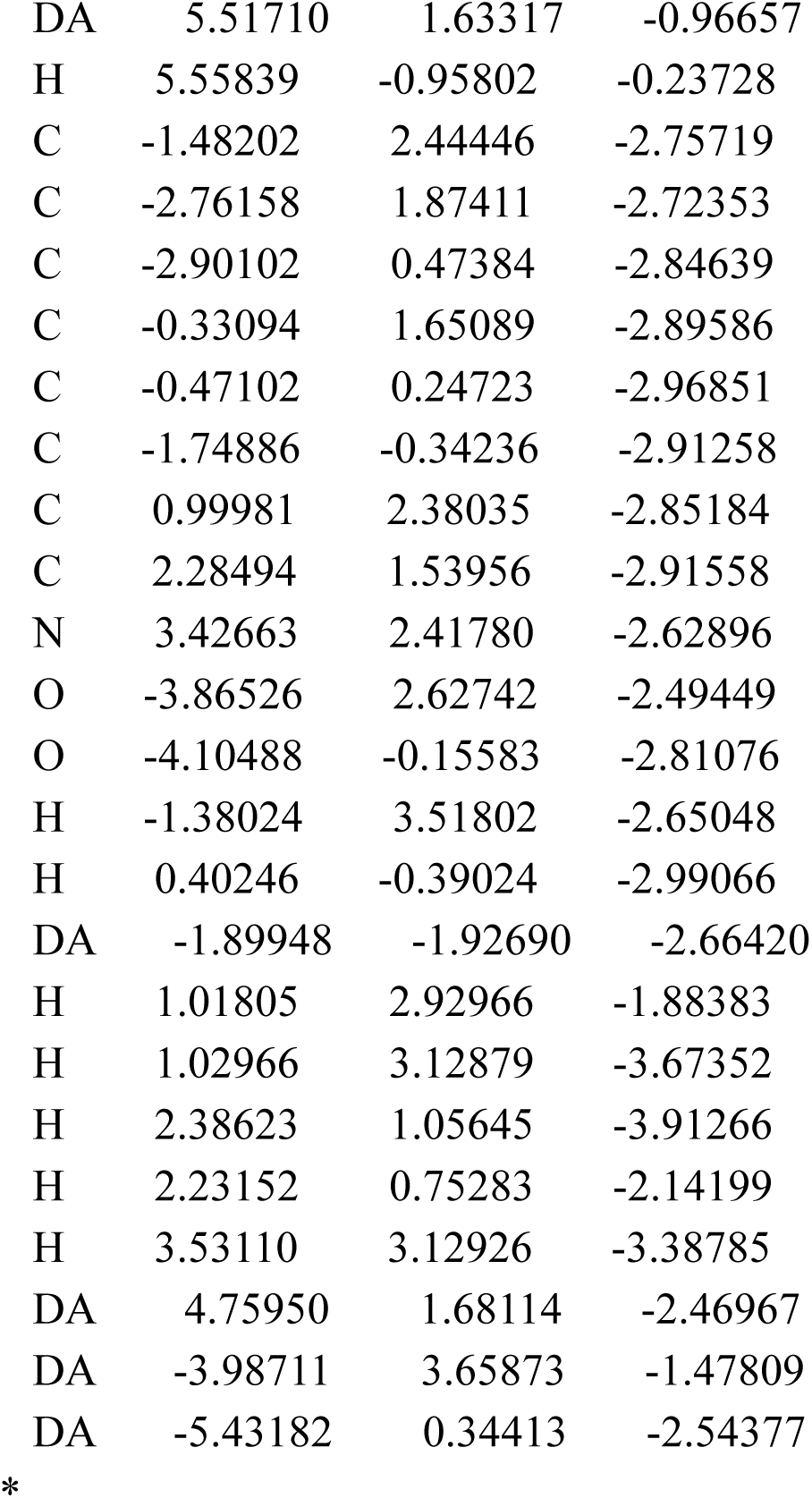

### Supplemental information 4: Molecular input for dopamine

|  |  |  |  |
| --- | --- | --- | --- |
| C | 0.71748 | 0.05390 | 1.00286 |
| C | -0.66792 | -0.12239 | 1.05882 |
| C | -1.40519 | -0.13296 | -0.12039 |
| C | -0.76352 | 0.02830 | -1.34596 |
| C | 0.62226 | 0.20572 | -1.40463 |
| C | 1.38368 | 0.21200 | -0.22546 |
| O | -1.50818 | 0.02223 | -2.49396 |
| O | -2.76243 | -0.29169 | -0.08492 |
| C | 2.87894 | 0.45877 | -0.20113 |
| C | 3.66256 | 0.05828 | -1.45828 |
| N | 5.09891 | 0.25640 | -1.26069 |
| H | 1.26180 | 0.06832 | 1.87952 |
| H | -1.14022 | -0.24380 | 1.96834 |
| H | 1.08654 | 0.33242 | -2.31754 |
| H | -2.43650 | -0.09700 | -2.44961 |
| H | -3.20008 | -0.40064 | 0.73639 |
| H | 3.28502 | -0.08985 | 0.62289 |
| H | 2.98820 | 1.52061 | -0.12725 |
| H | 3.33564 | 0.65862 | -2.28145 |
| H | 3.48364 | -0.97752 | -1.65833 |
| H | 5.28308 | 1.24113 | -1.06890 |
| H | 5.60007 | -0.02542 | -2.10320 |
| H | 5.41475 | -0.30842 | -0.47226 |

\*

### Supplemental information 5: MOlecular input for dopamine binding on carbon surface capped with hydrogens

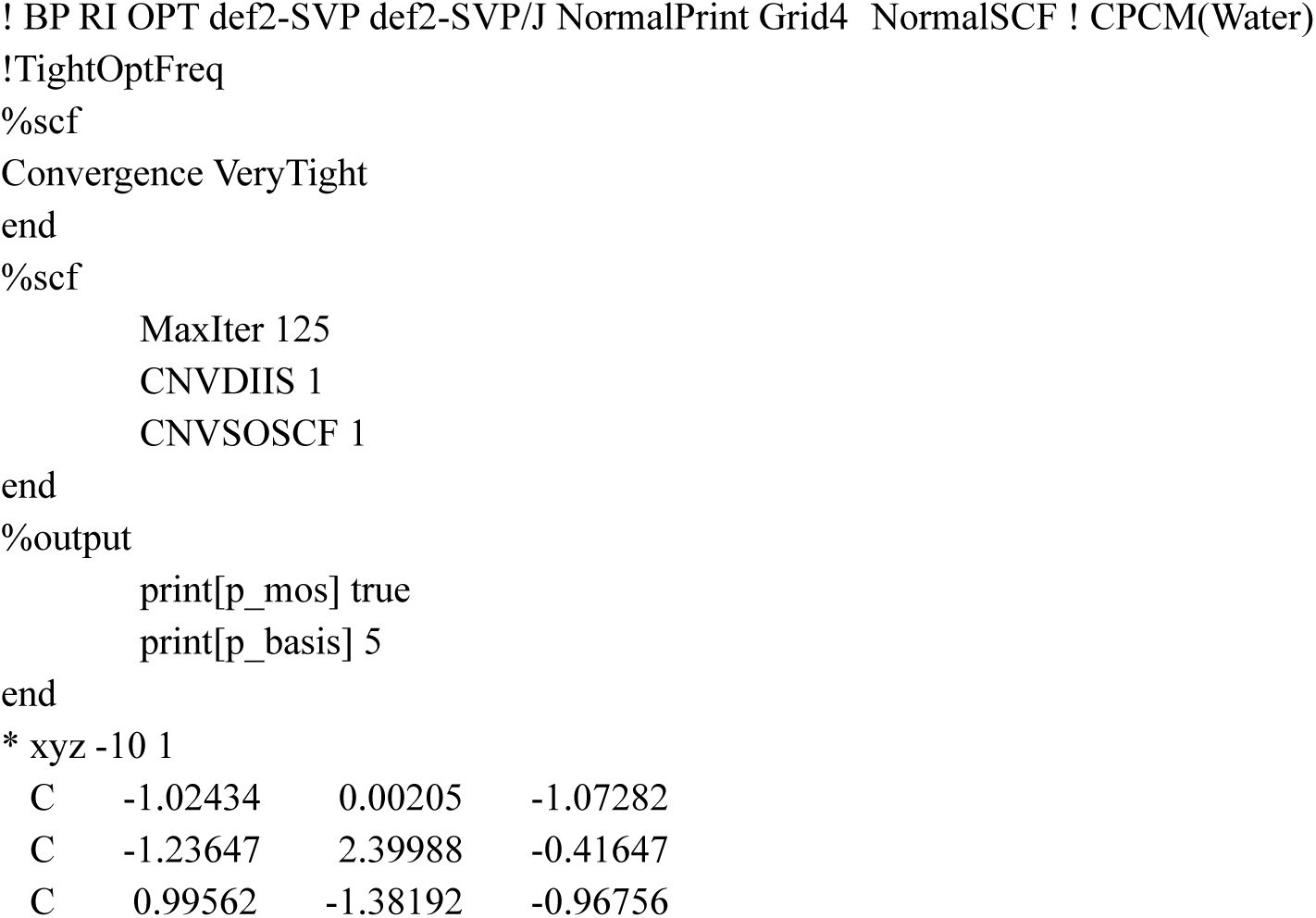
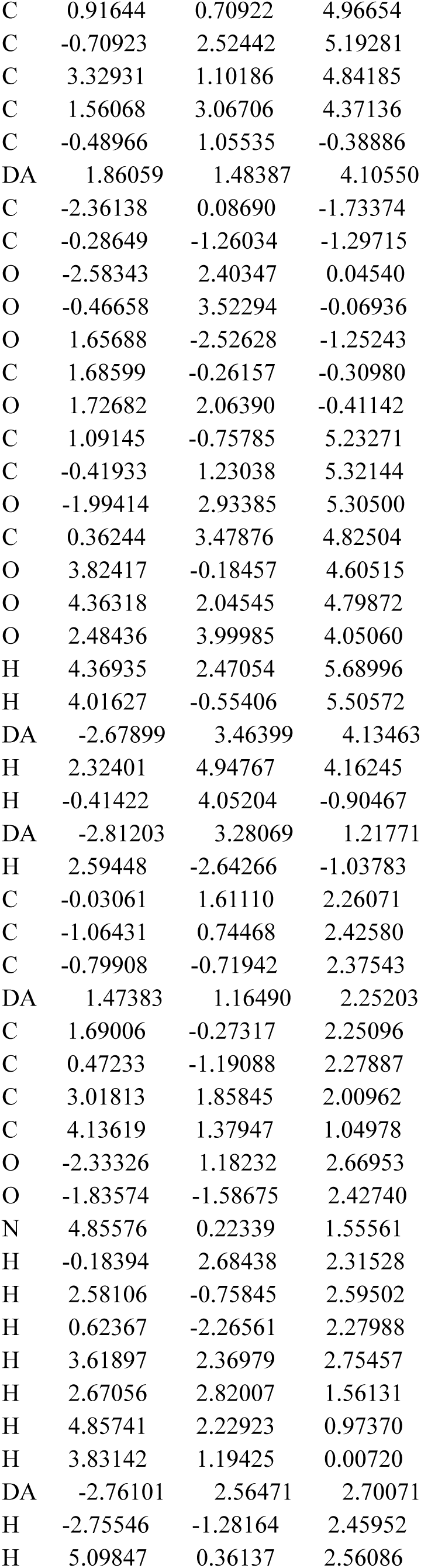
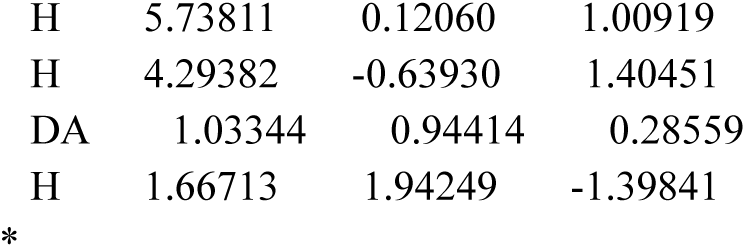

### Supplemental information 6: Molecular input for dopamine binding to N-dope graphite

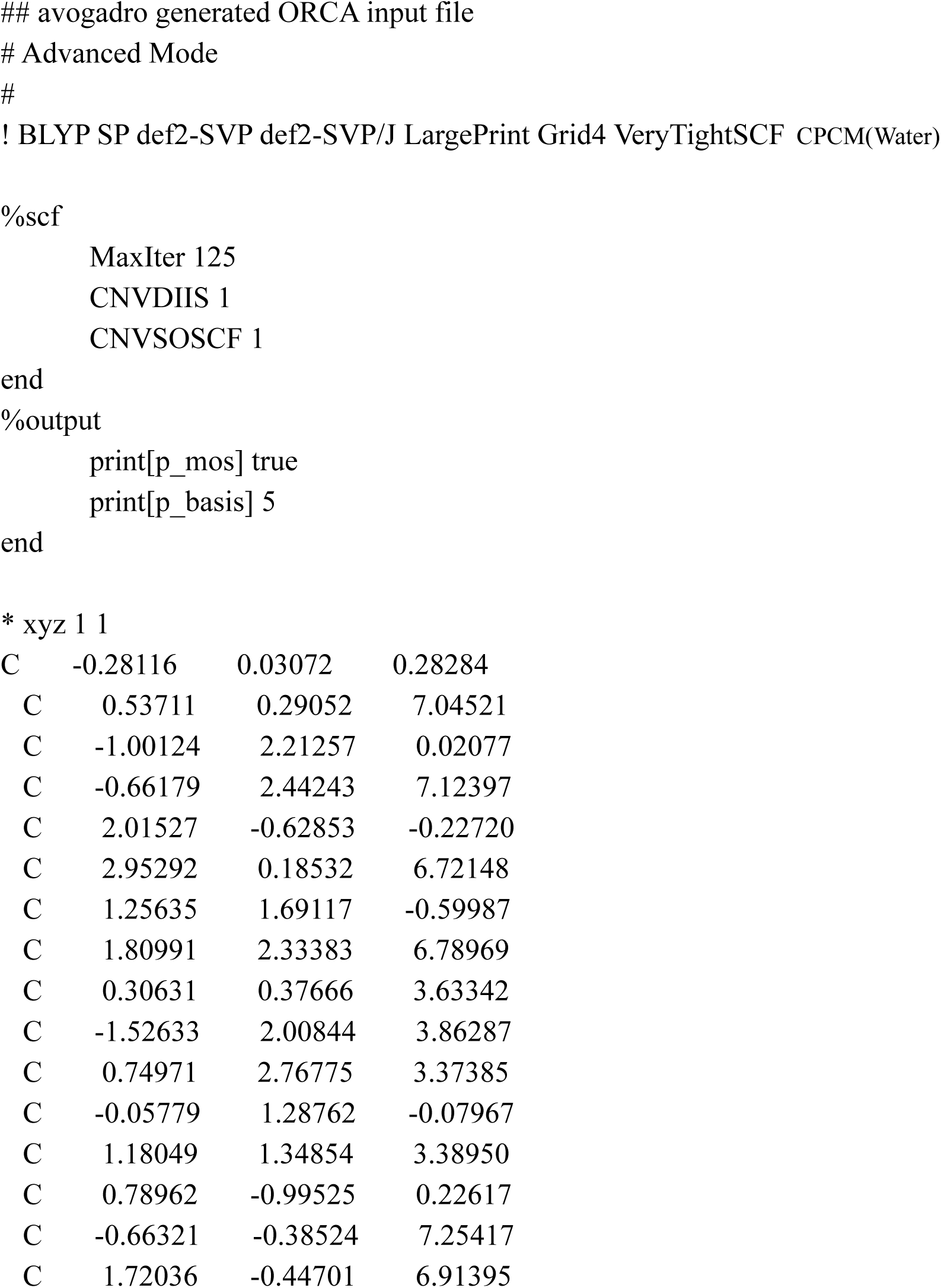
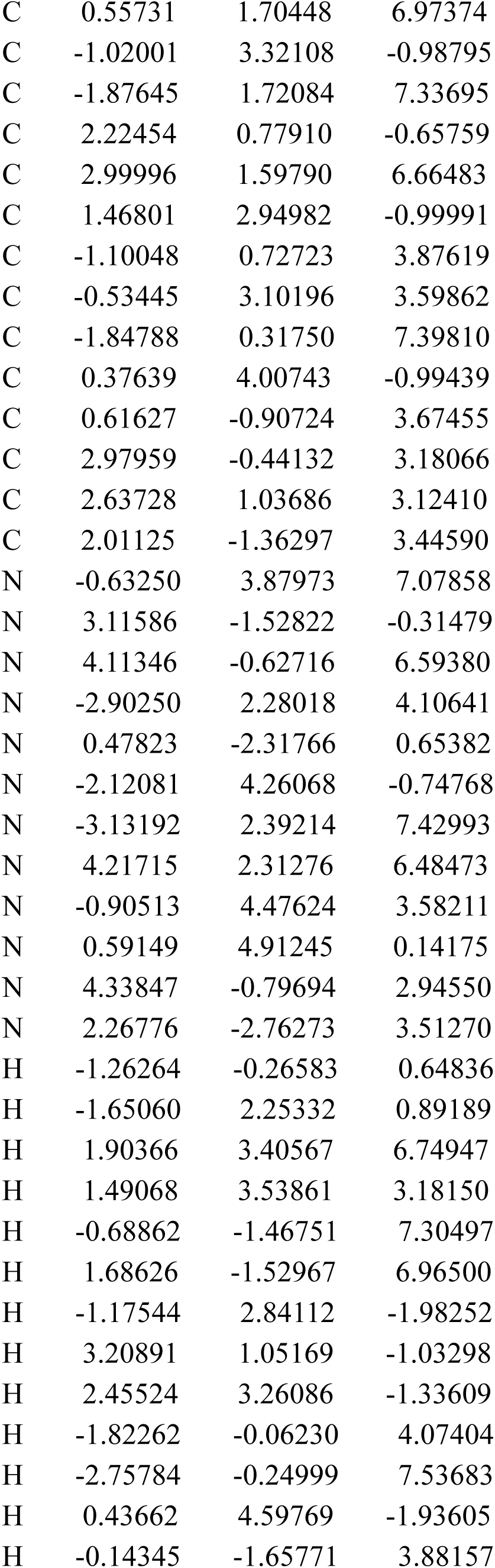
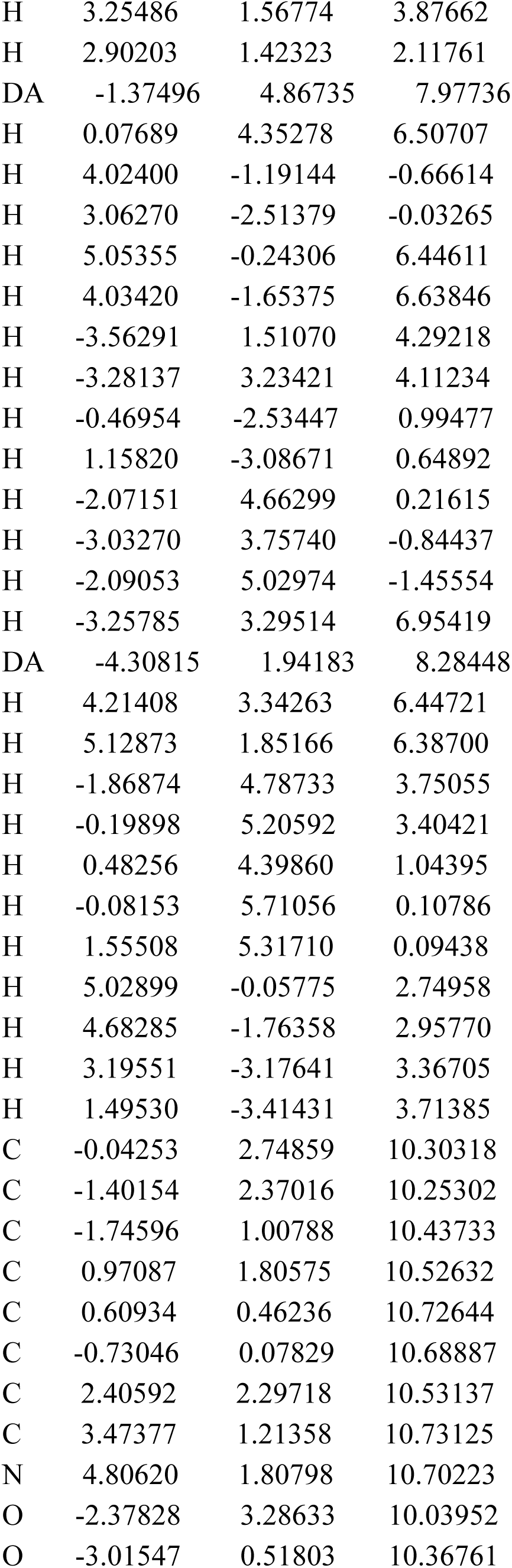
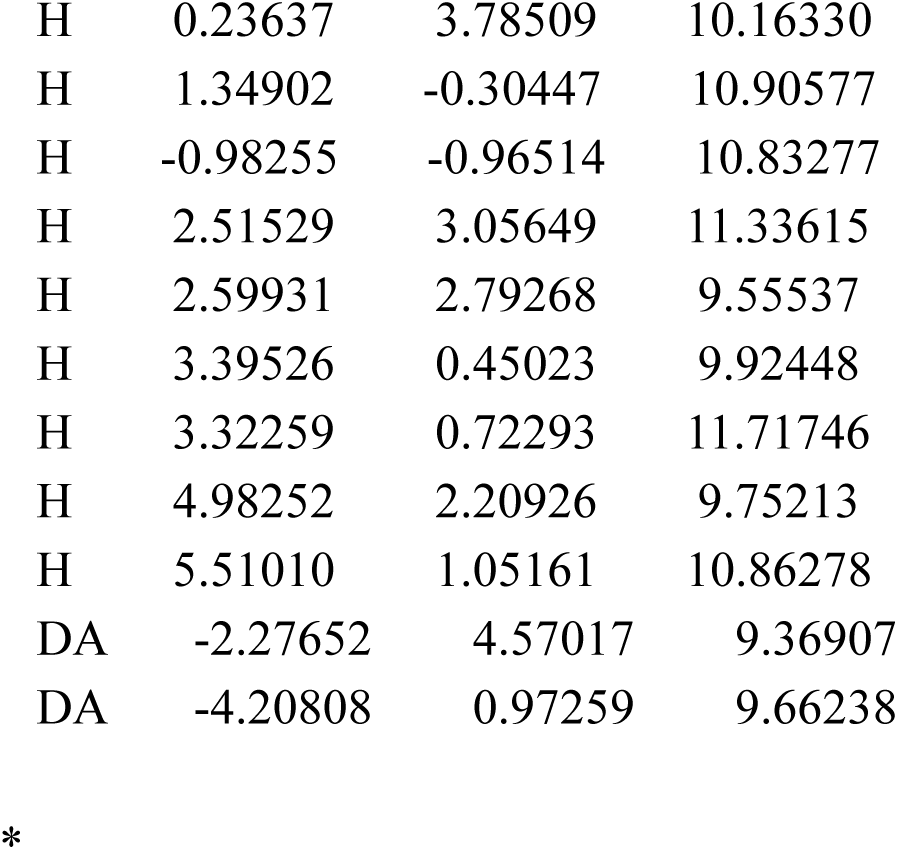

|  |  |  |  |
| --- | --- | --- | --- |
| C | -0.28116 | 0.03072 | 0.28284 |
| C | 0.53711 | 0.29052 | 7.04521 |
| C | -1.00124 | 2.21257 | 0.02077 |
| C | -0.66179 | 2.44243 | 7.12397 |
| C | 2.01527 | -0.62853 | -0.22720 |
| C | 2.95292 | 0.18532 | 6.72148 |
| C | 1.25635 | 1.69117 | -0.59987 |
| C | 1.80991 | 2.33383 | 6.78969 |
| C | 0.30631 | 0.37666 | 3.63342 |
| C | -1.52633 | 2.00844 | 3.86287 |
| C | 0.74971 | 2.76775 | 3.37385 |
| C | -0.05779 | 1.28762 | -0.07967 |
| C | 1.18049 | 1.34854 | 3.38950 |
| C | 0.78962 | -0.99525 | 0.22617 |
| C | -0.66321 | -0.38524 | 7.25417 |
| C | 1.72036 | -0.44701 | 6.91395 |
| C | 0.55731 | 1.70448 | 6.97374 |
| C | -1.02001 | 3.32108 | -0.98795 |
| C | -1.87645 | 1.72084 | 7.33695 |
| C | 2.22454 | 0.77910 | -0.65759 |
| C | 2.99996 | 1.59790 | 6.66483 |
| C | 1.46801 | 2.94982 | -0.99991 |
| C | -1.10048 | 0.72723 | 3.87619 |
| C | -0.53445 | 3.10196 | 3.59862 |
| C | -1.84788 | 0.31750 | 7.39810 |
| C | 0.37639 | 4.00743 | -0.99439 |
| C | 0.61627 | -0.90724 | 3.67455 |
| C | 2.97959 | -0.44132 | 3.18066 |
| C | 2.63728 | 1.03686 | 3.12410 |
| C | 2.01125 | -1.36297 | 3.44590 |
| N | -0.63250 | 3.87973 | 7.07858 |
| N | 3.11586 | -1.52822 | -0.31479 |
| N | 4.11346 | -0.62716 | 6.59380 |
| N | -2.90250 | 2.28018 | 4.10641 |
| N | 0.47823 | -2.31766 | 0.65382 |
| N | -2.12081 | 4.26068 | -0.74768 |
| N | -3.13192 | 2.39214 | 7.42993 |
| N | 4.21715 | 2.31276 | 6.48473 |
| N | -0.90513 | 4.47624 | 3.58211 |
| N | 0.59149 | 4.91245 | 0.14175 |
| N | 4.33847 | -0.79694 | 2.94550 |
| N | 2.26776 | -2.76273 | 3.51270 |
| H | -1.26264 | -0.26583 | 0.64836 |
| H | -1.65060 | 2.25332 | 0.89189 |
| H | 1.90366 | 3.40567 | 6.74947 |
| H | 1.49068 | 3.53861 | 3.18150 |
| H | -0.68862 | -1.46751 | 7.30497 |
| H | 1.68626 | -1.52967 | 6.96500 |
| H | -1.17544 | 2.84112 | -1.98252 |
| H | 3.20891 | 1.05169 | -1.03298 |
| H | 2.45524 | 3.26086 | -1.33609 |
| H | -1.82262 | -0.06230 | 4.07404 |
| H | -2.75784 | -0.24999 | 7.53683 |
| H | 0.43662 | 4.59769 | -1.93605 |
| H | -0.14345 | -1.65771 | 3.88157 |
| H | 3.25486 | 1.56774 | 3.87662 |
| H | 2.90203 | 1.42323 | 2.11761 |
| DA | -1.37496 | 4.86735 | 7.97736 |
| H | 0.07689 | 4.35278 | 6.50707 |
| H | 4.02400 | -1.19144 | -0.66614 |
| H | 3.06270 | -2.51379 | -0.03265 |
| H | 5.05355 | -0.24306 | 6.44611 |
| H | 4.03420 | -1.65375 | 6.63846 |
| H | -3.56291 | 1.51070 | 4.29218 |
| H | -3.28137 | 3.23421 | 4.11234 |
| H | -0.46954 | -2.53447 | 0.99477 |
| H | 1.15820 | -3.08671 | 0.64892 |
| H | -2.07151 | 4.66299 | 0.21615 |
| H | -3.03270 | 3.75740 | -0.84437 |
| H | -2.09053 | 5.02974 | -1.45554 |
| H | -3.25785 | 3.29514 | 6.95419 |
| DA | -4.30815 | 1.94183 | 8.28448 |
| H | 4.21408 | 3.34263 | 6.44721 |
| H | 5.12873 | 1.85166 | 6.38700 |
| H | -1.86874 | 4.78733 | 3.75055 |
| H | -0.19898 | 5.20592 | 3.40421 |
| H | 0.48256 | 4.39860 | 1.04395 |
| H | -0.08153 | 5.71056 | 0.10786 |
| H | 1.55508 | 5.31710 | 0.09438 |
| H | 5.02899 | -0.05775 | 2.74958 |
| H | 4.68285 | -1.76358 | 2.95770 |
| H | 3.19551 | -3.17641 | 3.36705 |
| H | 1.49530 | -3.41431 | 3.71385 |
| C | -0.04253 | 2.74859 | 10.30318 |
| C | -1.40154 | 2.37016 | 10.25302 |
| C | -1.74596 | 1.00788 | 10.43733 |
| C | 0.97087 | 1.80575 | 10.52632 |
| C | 0.60934 | 0.46236 | 10.72644 |
| C | -0.73046 | 0.07829 | 10.68887 |
| C | 2.40592 | 2.29718 | 10.53137 |
| C | 3.47377 | 1.21358 | 10.73125 |
| N | 4.80620 | 1.80798 | 10.70223 |
| O | -2.37828 | 3.28633 | 10.03952 |
| O | -3.01547 | 0.51803 | 10.36761 |
| H | 0.23637 | 3.78509 | 10.16330 |
| H | 1.34902 | -0.30447 | 10.90577 |
| H | -0.98255 | -0.96514 | 10.83277 |
| H | 2.51529 | 3.05649 | 11.33615 |
| H | 2.59931 | 2.79268 | 9.55537 |
| H | 3.39526 | 0.45023 | 9.92448 |
| H | 3.32259 | 0.72293 | 11.71746 |
| H | 4.98252 | 2.20926 | 9.75213 |
| H | 5.51010 | 1.05161 | 10.86278 |
| DA | -2.27652 | 4.57017 | 9.36907 |
| DA | -4.20808 | 0.97259 | 9.66238 |

\*

### Supplemental information 7: Molecular input for serotonin

|  |  |  |  |
| --- | --- | --- | --- |
| C | 0.47918 | 0.16604 | 2.11923 |
| C | -0.84655 | 0.46338 | 1.73327 |
| C | -1.21537 | 0.39631 | 0.38076 |
| H | -2.23106 | 0.62312 | 0.07969 |
| C | -0.28341 | 0.03817 | -0.59186 |
| H | -0.58602 | -0.00826 | -1.63047 |
| C | 1.03280 | -0.25838 | -0.22499 |
| O | 1.94553 | -0.60600 | -1.15482 |
| C | 1.41014 | -0.19482 | 1.11991 |
| H | 2.43594 | -0.43081 | 1.36633 |
| C | -1.89538 | 0.85638 | 2.73889 |
| C | 0.87043 | 0.23692 | 3.56182 |
| C | -0.06407 | 0.58939 | 4.47142 |
| C | 2.30914 | -0.09131 | 3.96465 |
| C | 2.64379 | -0.00046 | 5.46186 |
| N | 4.04792 | -0.32974 | 5.68570 |
| N | -1.40183 | 0.89282 | 4.09950 |
| H | 1.72671 | -0.66199 | -2.09603 |
| H | -2.73264 | 0.12778 | 2.68584 |
| H | -2.28251 | 1.86422 | 2.47616 |
| H | 0.17369 | 0.65367 | 5.52578 |
| H | 2.98542 | 0.60867 | 3.42741 |
| H | 2.52746 | -1.12931 | 3.63199 |
| H | 2.46225 | 1.03708 | 5.81820 |
| H | 2.00551 | -0.70590 | 6.03945 |
| H | 4.25196 | -0.23701 | 6.70706 |
| H | 4.21493 | -1.32680 | 5.41728 |
| H | -2.06352 | 1.15457 | 4.84509 |

\*

### Supplemental information 8: Molecular input for serotonin binding with hydrogenated graphite

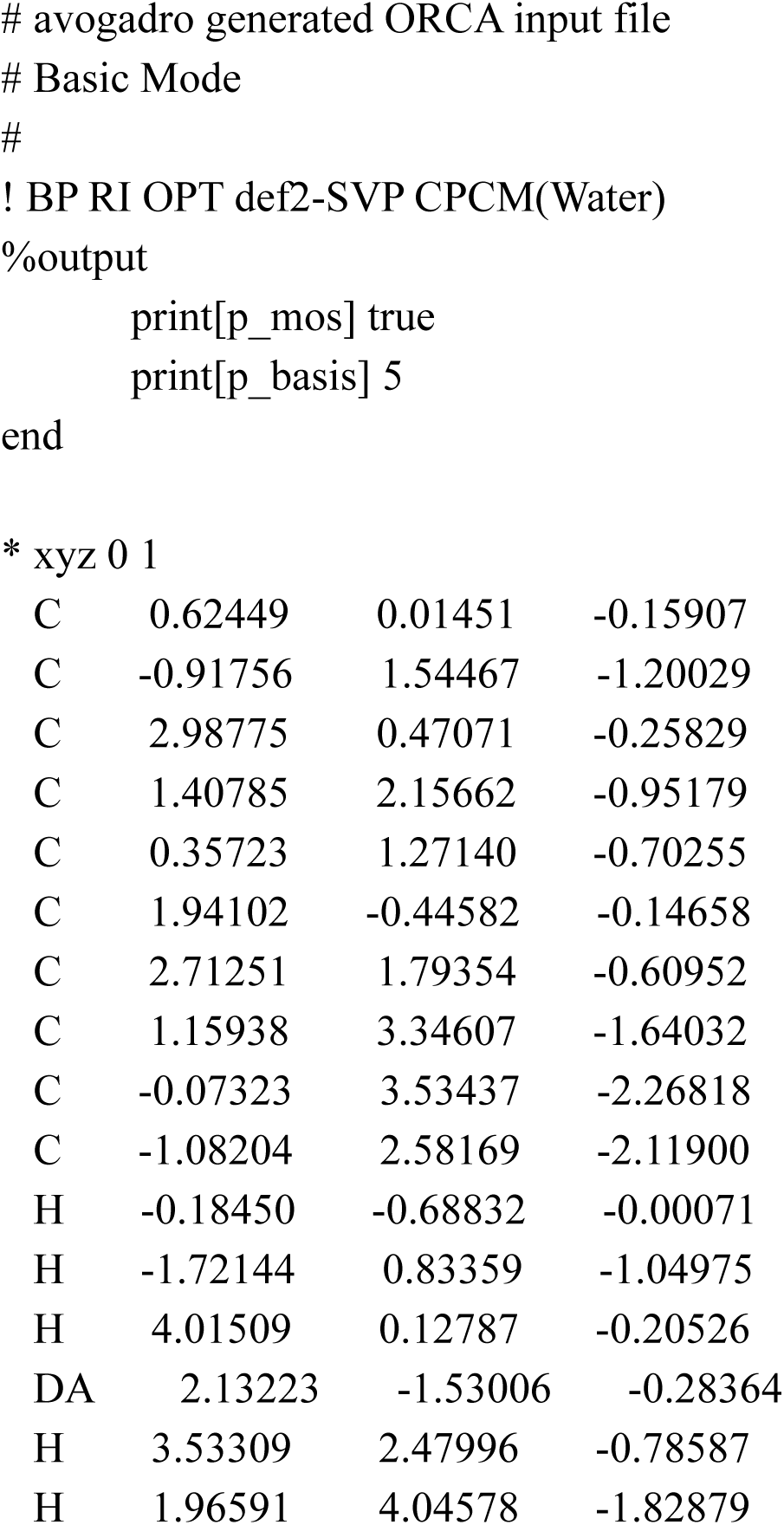
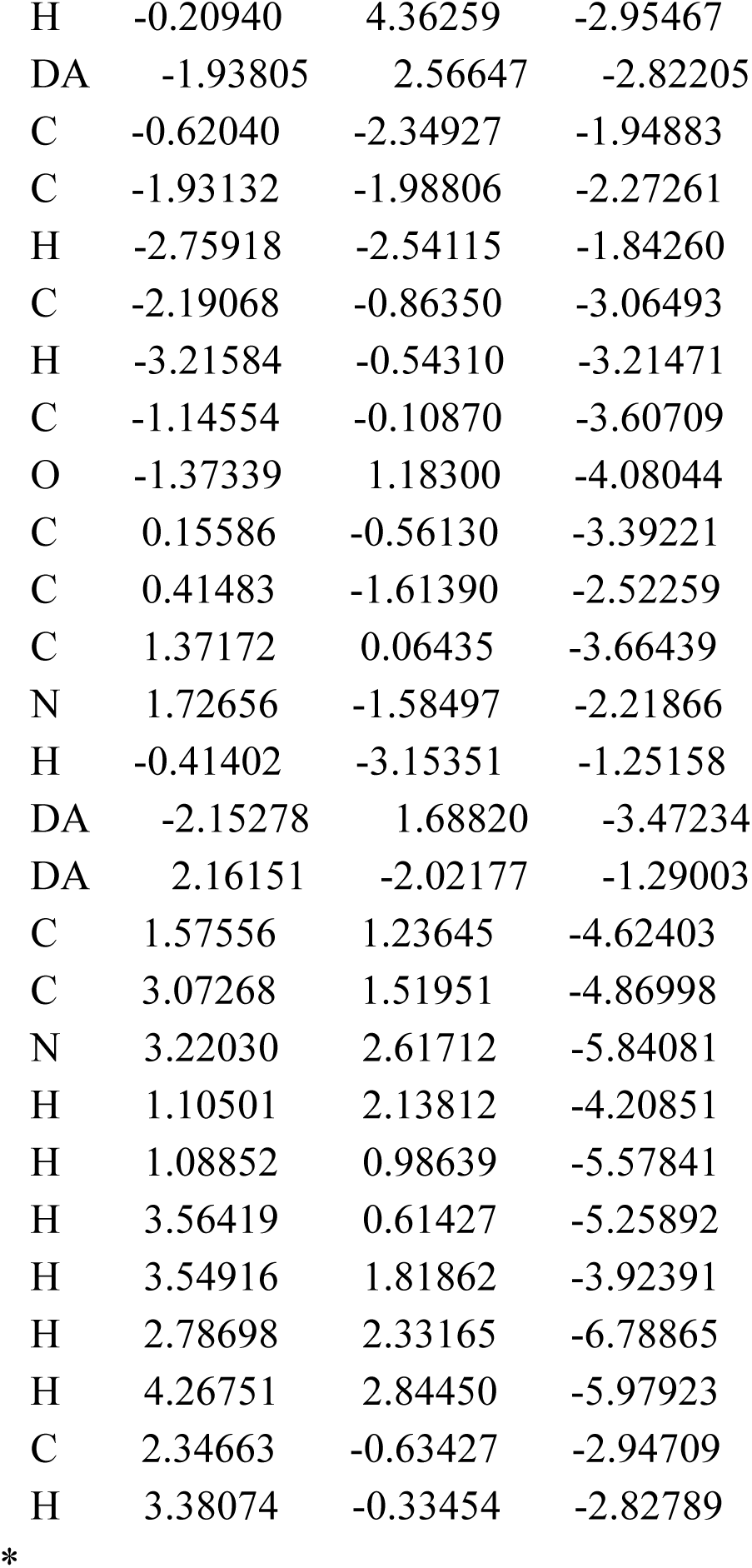

### Supplemental information 9 Molecular input for Serotonin binding with N-doped hydrogenated graphite

|  |  |  |  |
| --- | --- | --- | --- |
| C | -0.31326 | 0.12208 | 7.91322 |
| C | -1.56800 | 2.04973 | 7.17856 |
| C | 1.63621 | 0.31630 | 9.34873 |
| C | 0.29346 | 2.23588 | 8.67586 |
| C | 0.03068 | 3.51040 | 2.06366 |
| C | -0.96266 | 4.16639 | 4.17596 |
| C | 2.03030 | 2.35824 | 1.96200 |
| C | 1.13082 | 2.87939 | 4.14387 |
| C | 1.05975 | 2.92054 | 2.67056 |
| C | -1.18561 | -0.71538 | 7.07852 |
| C | 0.66862 | -0.50027 | 8.56684 |
| C | -0.48874 | 1.42835 | 7.95916 |
| C | -2.34871 | 1.30420 | 6.37898 |
| C | 1.45367 | 1.64833 | 9.40243 |
| C | -1.04216 | 4.15709 | 2.84300 |
| C | 0.18958 | 3.51415 | 4.83980 |
| C | 3.10591 | 1.59860 | 2.61467 |
| C | 2.11891 | 2.20761 | 4.74601 |
| C | -2.12616 | -0.15874 | 6.28955 |
| C | 3.14633 | 1.48491 | 3.95476 |
| H | -1.73192 | 3.12384 | 7.21518 |
| H | 2.47211 | -0.15247 | 9.85842 |
| H | 0.14378 | 3.31206 | 8.69880 |
| H | -0.06357 | 3.49948 | 0.97936 |
| H | -1.73820 | 4.64233 | 4.76694 |
| H | 2.02535 | 2.38936 | 0.87439 |
| H | -1.04473 | -1.78835 | 7.07768 |
| H | 0.81298 | -1.57623 | 8.50626 |
| H | -3.12498 | 1.78221 | 5.78854 |
| H | 2.13990 | 2.28032 | 9.95698 |
| H | -1.88176 | 4.62076 | 2.33470 |
| H | 0.24233 | 3.55586 | 5.92140 |
| H | 3.83969 | 1.12403 | 1.97807 |
| H | 2.15508 | 2.13585 | 5.83174 |
| DA | -2.96621 | -0.99426 | 5.23747 |
| DA | 4.23609 | 0.55941 | 4.65813 |
| H | 2.36803 | -1.60912 | 5.28885 |
| C | 1.80221 | -1.76898 | 4.38055 |
| C | 0.45252 | -2.09958 | 4.44860 |
| H | 0.02980 | -2.20850 | 5.40912 |
| C | -0.33483 | -2.29397 | 3.29090 |
| O | -1.66873 | -2.56994 | 3.37354 |
| C | 0.29335 | -2.20483 | 2.02297 |
| H | -0.27254 | -2.35492 | 1.11217 |
| C | 1.66511 | -1.90210 | 1.97849 |
| C | 2.38709 | -1.67179 | 3.12386 |
| C | 2.54507 | -1.72461 | 0.92062 |
| C | 3.77874 | -1.40217 | 1.49386 |
| N | 3.66031 | -1.34063 | 2.83659 |
| C | 2.21382 | -1.79651 | -0.54624 |
| C | 3.40990 | -1.42379 | -1.43423 |
| N | 3.02943 | -1.43359 | -2.84113 |
| DA | -2.52998 | -2.47680 | 4.55565 |
| H | 4.68790 | -1.15388 | 0.96272 |
| DA | 4.72423 | -0.81384 | 3.80766 |
| H | 1.39465 | -1.07196 | -0.74674 |
| H | 1.85769 | -2.81887 | -0.79226 |
| H | 4.24813 | -2.13697 | -1.26736 |
| H | 3.75343 | -0.39805 | -1.17003 |
| H | 2.78572 | -2.41025 | -3.12254 |
| H | 3.85712 | -1.13311 | -3.40588 |

\*

### Supplemental information 10: Dopamine molecular input binding with graphite doped with oxygen

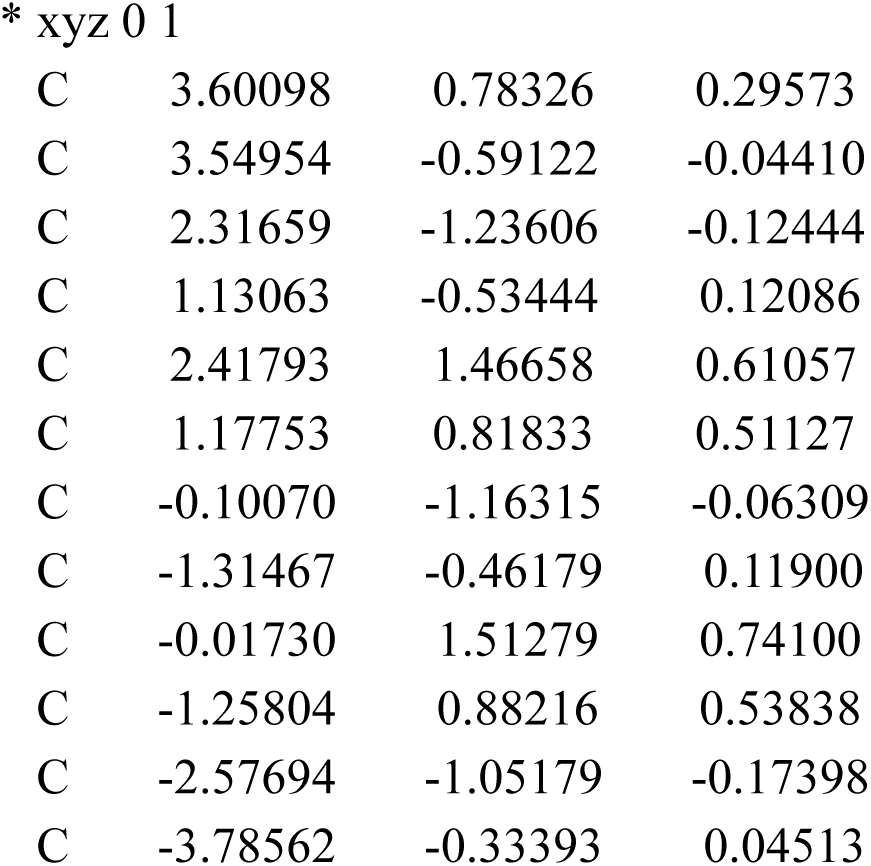
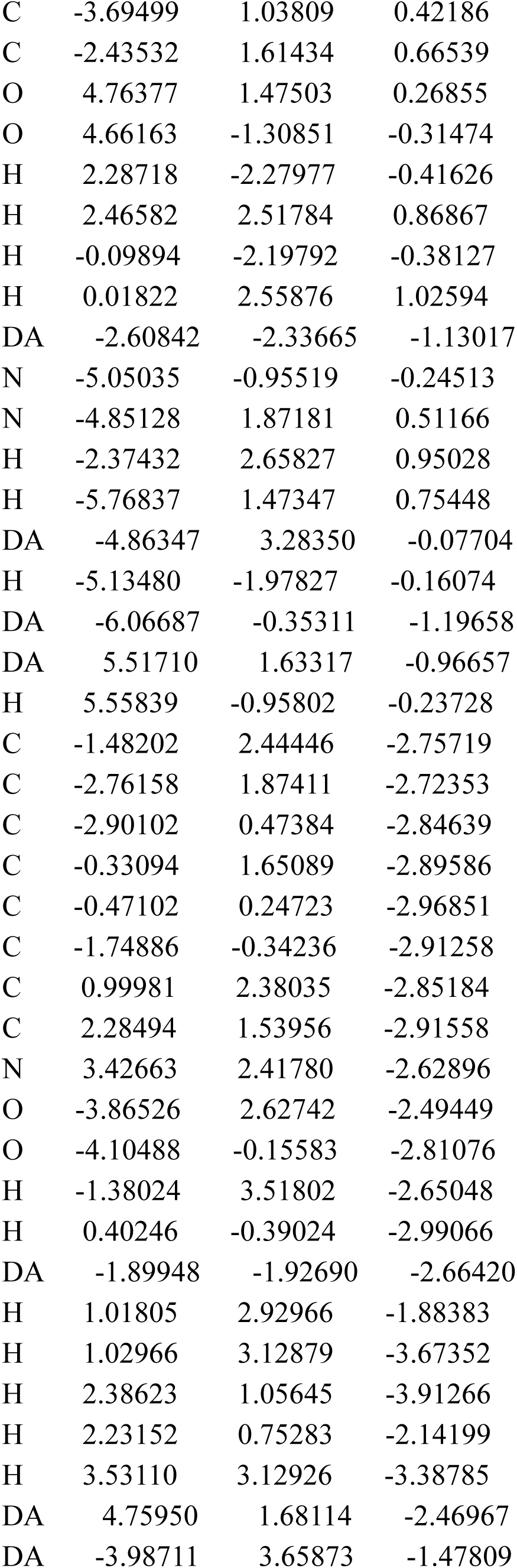
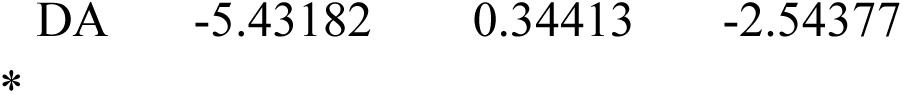

